# *Treponema pallidum* eight-stranded beta-barrels: an evolving, versatile family of outer membrane proteins in the service of stealth pathogenicity

**DOI:** 10.64898/2026.11.15.766778

**Authors:** Everton B. Bettin, Farhang Aghakhanian, B. Dillon Vannest, André A. Grassmann, Jonathan B. Parr, Kelly L. Hawley, Justin D. Radolf, Melissa J. Caimano

**Affiliations:** Department of Medicine, UConn Health, Farmington, Connecticut, USA; University of North Carolina at Chapel Hill, Chapel Hill, North Carolina, USA; Department of Pediatrics, UConn Health, Farmington, Connecticut, USA; Department of Molecular Biology and Biophysics, UConn Health, Farmington, Connecticut, USA; Connecticut Children’s Research Institute, Hartford, Connecticut, USA; Divisions of Research and Infectious Diseases, Connecticut Children’s, Hartford, Connecticut, USA

**Author notes:** Corresponding author: Melissa J. Caimano, Ph.D. Department of Medicine UConn Health, 263 Farmington Avenue, Farmington, CT 06030-3715.

**Keywords:** *Treponema pallidum*, syphilis, outer membrane proteins, beta-barrel proteins, DUF2715, OMP/PagP_b-brl

## Abstract

Syphilis is a multistage sexually transmitted infection caused by the highly invasive and immunoevasive spirochete *Treponema pallidum* (*TPA*). *TPA* depends on the import of many host-derived nutrients to sustain its auxotrophic lifestyle. Moreover, its invasive nature requires interactions with diverse host molecules to successfully disseminate and persist. The outer membrane proteins (OMPs) mediating these processes remain largely uncharacterized. In other bacteria, 8-stranded β-barrels (8SβBs) perform such functions, including nutrient uptake and host interactions. Here, we investigated the composition, structural diversity and evolutionary dynamics of the 8SβB family in *TPA*. Reexamination of the *TPA* OMPeome revealed four previously unrecognized members in addition to the four known 8SβBs, expanding the family to eight members. Based on RNAseq, all eight were transcribed in treponemes harvested from rabbit testes. Predicted structures revealed substantial differences in channel properties and extracellular loop (ECL) architecture, consistent with functional diversification. AlphaFold 3 modeling predicted interactions with diverse host molecules implicated in nutrient uptake, adherence, and immune regulation, providing testable hypotheses for the functional specialization of individual 8SβBs. Despite containing predicted B-cell epitopes, most *TPA* 8SβB ECLs exhibited limited antigenicity during rabbit infection. Analysis of global clinical strains revealed distinct patterns of sequence variation across the 8SβB family, although most were highly conserved and appeared to be under purifying selection. Variation within hypermutable intragenic polyguanosine tracts emerged as an additional potential mechanism for modulating 8SβB repertoire composition. Our findings highlight the 8SβB family as an important component of the molecular architecture underlying *TPA* stealth pathogenicity and reveal an ongoing mechanism through which the spirochete diversifies its OMP repertoire within circulating populations.

**Author summary:** Syphilis is a sexually transmitted infection caused by *Treponema pallidum* subspecies *pallidum* (*TPA*). To survive and replicate within its obligate human host, *TPA* must import host-derived nutrients across its outer membrane. The *TPA* outer membrane also serves as a platform for surface molecules that function at the host-pathogen interface. By leveraging advances in bioinformatics and structural modeling, we have made significant strides in uncovering the repertoire of outer membrane proteins (OMPs) that enable syphilis spirochetes to infect and persist in humans. Our Herein, we describe four previously unrecognized eight-stranded beta-barrel (8SβB) paralogs. Despite their diminutive size, 8SβBs serve a surprisingly broad range of physiologic and virulence-related functions. Based on their predicted structural and biophysical properties, the eight members of this expanded paralogous family likely serve overlapping but nonredundant roles in *TPA*, including nutrient uptake and host-pathogen interactions. Analysis of clinical strains from diverse geographic sites suggests that variability in intragenic polyG tracts within 8SβBs reshapes and expands this OMP family on a global scale. Collectively, our findings revealed a seemingly versatile OMP family whose functional specialization and expansion enable *TPA* to meet the demands of stealth pathogenicity in individuals and across populations.

## Introduction

Syphilis is a multi-stage, sexually transmitted infection caused by the highly invasive and immunoevasive spirochete *Treponema pallidum* subspecies (subsp.) *pallidum* (*TPA*) [1, 2]. Following transmission, *TPA* disseminates rapidly from the site of inoculation, infecting diverse organs, including the cardiovascular and central nervous systems [3]. *TPA* is also one of the few bacterial pathogens capable of crossing the placental barrier during pregnancy, often leading to stillbirth or other adverse birth outcomes [3, 4]. In the United States, more than 200,000 new cases of syphilis were reported in 2023, the highest number since the 1950s [5]. According to the World Health Organization, ∼7-8 million adults acquire venereal syphilis annually worldwide, while the incidence of congenital syphilis has surged globally to ∼700,000 cases per year [6]. Deconvoluting the virulence properties that enable *TPA* to adapt so successfully to geographically and demographically diverse populations poses a formidable challenge to syphilis investigators and their efforts to curtail the spread of its causative agent.

To survive and replicate within its obligate human host, *TPA*, an extreme auxotroph, must import an array of host-derived nutrients across its outer membrane. The *TPA* outer membrane (OM) also serves as a platform for surface molecules that function at the host-pathogen interface [2, 7]. By leveraging recent advances in bioinformatics and structural modeling, we have made significant strides in uncovering the repertoire of outer membrane proteins (OMPs) that enable syphilis spirochetes to meet the physiologic and virulence-related requirements for survival and persistence within their obligate human host [7, 8]. The *TPA* OMPeome includes BamA and LptD, involved in OM biogenesis, and four distinct paralogous OMP families presumably involved in importation of nutrients or extrusion of noxious substances across the OM: *TPA* repeat proteins (Tprs), OM factors for efflux pumps (OMFs), orthologs for FadL long-chain fatty acid transporters, and 8-stranded β-barrels (8SβBs) [7, 9]. The discovery that the spirochete’s genome encodes multiple paralogous families of OMPs helps resolve a central paradox of *TPA* cell biology, namely, how a diderm bacterium with an extraordinarily low density of integral OMPs can implement such a complex, systemic parasitic strategy [7, 10].

Despite their diminutive size, 8SβBs serve a surprisingly broad range of biological roles in bacteria [11], including the passive diffusion of small molecules across the OM (*e.g*., OmpW [12], CarO [13] and OprG [14]), stabilization of the OM protein lattice (*e.g*., OmpA [15]), and modification of lipid A by the addition of palmitic acid from a phospholipid donor (*e.g*., PagP [16]). 8SβBs also serve a variety of virulence-related functions. For example, Opacity-associated (Opa) proteins in *Neisseria* spp. bind members of the carcinoembryonic antigen-related cell adhesion molecule (CEACAM) family, promoting bacterial attachment and modulation of host immune responses [17]. The 8SβBs Ail and Rck in *Yersinia* and *Salmonella* spp. contribute to complement inhibition through well-characterized interactions with vitronectin [18] and complement regulatory factor H, respectively [19, 20].

In our initial assessment of the *TPA* Nichols OMPeome, we described four 8SβB paralogs [7]. Our reexamination of the *TPA* proteome using updated computational tools, described herein, brought to light four previously unrecognized paralogs. Based on their predicted structural and biophysical properties, the eight members of this expanded paralogous family likely serve overlapping but nonredundant roles during infection. RNA-seq confirmed that all eight are transcribed during rabbit infection and that they elicit mostly weak antibody responses directed against their extracellular loops (ECLs). Structural predictions further suggested that individual paralogs interact with CEACAMs, vitronectin, factor H, and cadherins. Analysis of *TPA* clinical strains from diverse geographic sites revealed that variability in intragenic polymeric guanosine (polyG) nucleotide tracts within the 8SβB-encoding regions reshapes and expands this OMP family globally. Collectively, these findings reveal a seemingly versatile family of OMPs whose functional specialization and expansion enable *TPA* to meet the demands of stealth pathogenicity in individuals and across populations.

## Results

### *Treponema pallidum* subsp. *pallidum* encodes a distinct repertoire of 8-stranded β-barrels with predicted differences in channel properties and ECL architecture

We previously identified four coding sequences (TP0126, TP0479, TP0698 and TP0733) in *TPA* Nichols that are predicted to form single-domain 8SβBs [7] (Table 1). As part of our ongoing assessment of the *TPA* OMPeome, we searched the Nichols reference genome and proteome for domains belonging to the membrane β-barrel (MBB) clan (CL0193), which encompasses β-barrel transmembrane proteins identified in Gram-negative bacteria [21]. This search identified four additional 8SβBs - TP0346, TP0347, TP0619 and TP0697 (Table 1). A search of the *TPA* SS14 genome and proteome revealed syntenic, highly conserved (99-100%) orthologs for all eight 8SβBs identified in *TPA* Nichols (Table 1). Two (TP0126 and TP0733) contain an OMP/PagP_b-brl (IPR011250) domain, shared by structurally and functionally diverse 8SβBs (*e.g.,* OmpA [22, 23], OmpX [24], OmpW [12], and PagP [16]. The remaining six contain instead the <u>D</u>omain of <u>U</u>nknown <u>F</u>unction DUF2715 (IPR024471) [25, 26] found almost exclusively within *Treponema* species. Translational start site annotations for each coding region were evaluated and, in some cases revised, based on predicted SPaseI cleavage sites, putative ribosome binding sites, and RNA-seq mapped read data (see below) (S1 and S2 Figs). Notably, *TPA* Nichols and SS14 reference strains also encode three ‘degenerate’ 8SβBs (designated TP0127*, TP0315* and TP0618*) whose open reading frames are truncated due to frameshift mutations within hypermutable polymeric guanosine nucleotide (polyG) tracts (discussed below).

**Table 1.**
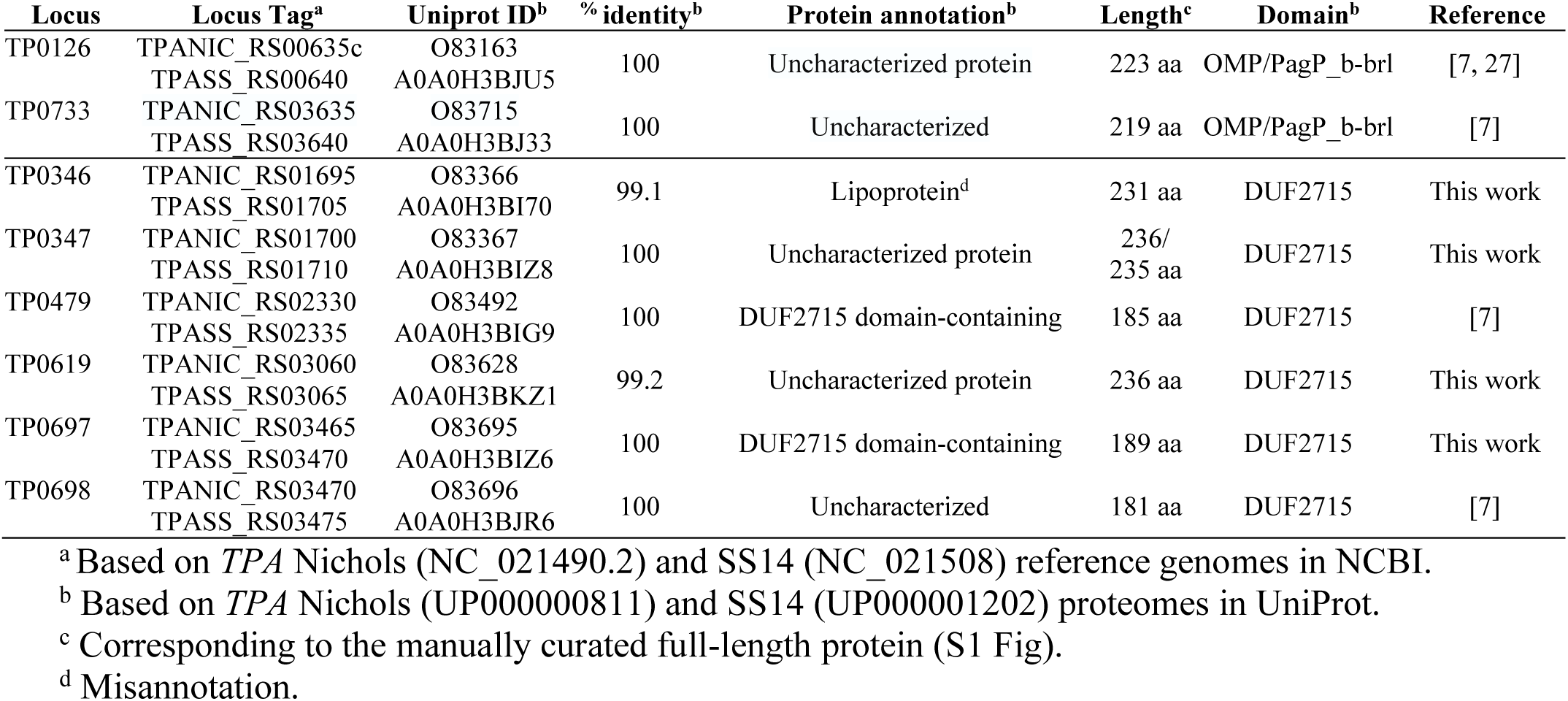
Full-length 8-stranded β-barrel proteins encoded by *TPA*.

| **Locus** | **Locus Tag<sup>a</sup>** | **Uniprot ID<sup>b</sup>** | **% identity<sup>b</sup>** | **Protein annotation<sup>b</sup>** | **Length<sup>c</sup>** | **Domain<sup>b</sup>** | **Reference** |
|---|---|---|---|---|---|---|---|
| TP0126 | TPANIC_RS00635c<br>TPASS_RS00640 | O83163<br>A0A0H3BJU5 | 100 | Uncharacterized protein | 223 aa | OMP/PagP_b-brl | [7, 27] |
| TP0733 | TPANIC_RS03635<br>TPASS_RS03640 | O83715<br>A0A0H3BJ33 | 100 | Uncharacterized | 219 aa | OMP/PagP_b-brl | [7] |
| TP0346 | TPANIC_RS01695<br>TPASS_RS01705 | O83366<br>A0A0H3BI70 | 99.1 | Lipoprotein<sup>d</sup> | 231 aa | DUF2715 | This work |
| TP0347 | TPANIC_RS01700<br>TPASS_RS01710 | O83367<br>A0A0H3BIZ8 | 100 | Uncharacterized protein | 236/235 aa | DUF2715 | This work |
| TP0479 | TPANIC_RS02330<br>TPASS_RS02335 | O83492<br>A0A0H3BIG9 | 100 | DUF2715 domain-containing | 185 aa | DUF2715 | [7] |
| TP0619 | TPANIC_RS03060<br>TPASS_RS03065 | O83628<br>A0A0H3BKZ1 | 99.2 | Uncharacterized protein | 236 aa | DUF2715 | This work |
| TP0697 | TPANIC_RS03465<br>TPASS_RS03470 | O83695<br>A0A0H3BIZ6 | 100 | DUF2715 domain-containing | 189 aa | DUF2715 | This work |
| TP0698 | TPANIC_RS03470<br>TPASS_RS03475 | O83696<br>A0A0H3BJR6 | 100 | Uncharacterized | 181 aa | DUF2715 | [7] |

<sup>a</sup> Based on *TPA* Nichols (NC_021490.2) and SS14 (NC_021508) reference genomes in NCBI.
<sup>b</sup> Based on *TPA* Nichols (UP000000811) and SS14 (UP000001202) proteomes in UniProt.
<sup>c</sup> Corresponding to the manually curated full-length protein (S1 Fig).
<sup>d</sup> Misannotation.

### Predicted differences in channel properties and ECL architecture suggest that *TPA* 8SβBs have diversified to support a broad range of functions

AlphaFold 3 (AF3) predicted that all eight mature proteins contain an 8SβB topology (Fig 1A). Like the well-characterized 8SβBs OmpA and OmpX from *E. coli* [28], the *TPA* orthologs are predicted to adopt a tilted orientation within the OM, guided by interactions of aromatic residues at the bilayer interface. TP0733 is unique in containing a 26-amino acid N-terminal periplasmic extension predicted to be intrinsically disordered (S3 and S4 Figs). Hydrophobicity analysis further confirmed that all eight *TPA* 8SβBs display membrane-spanning hydrophobic belts surrounding hydrophilic barrel interiors (S4 Fig). Based on MOLEonline [29], TP0126 and TP0733 form transmembrane channels with distinct electrostatic properties; the channel for TP0126 is weakly positive, while the TP0733 channel is strongly negative (Fig 1B). Spirochetes utilize ornithine, a positively charged amino acid, as a primary structural component of their peptidoglycan (PG) [30], but how *TPA* acquires this essential nutrient is unclear. In *Acinetobacter baumannii*, the import of ornithine is mediated by the 8SβB CarO [31]. We, therefore, hypothesized that TP0733 could contribute to ornithine uptake. Structure-based docking analysis predicted three potential ornithine-binding sites along the TP0733 β-barrel with predicted Vina scores ranging from −4.9 to −5.6 kcal/mol (Fig 1C). Predicted binding sites included multiple polar and charged residues capable of interacting with the zwitterionic ornithine molecule.

**Fig 1.**
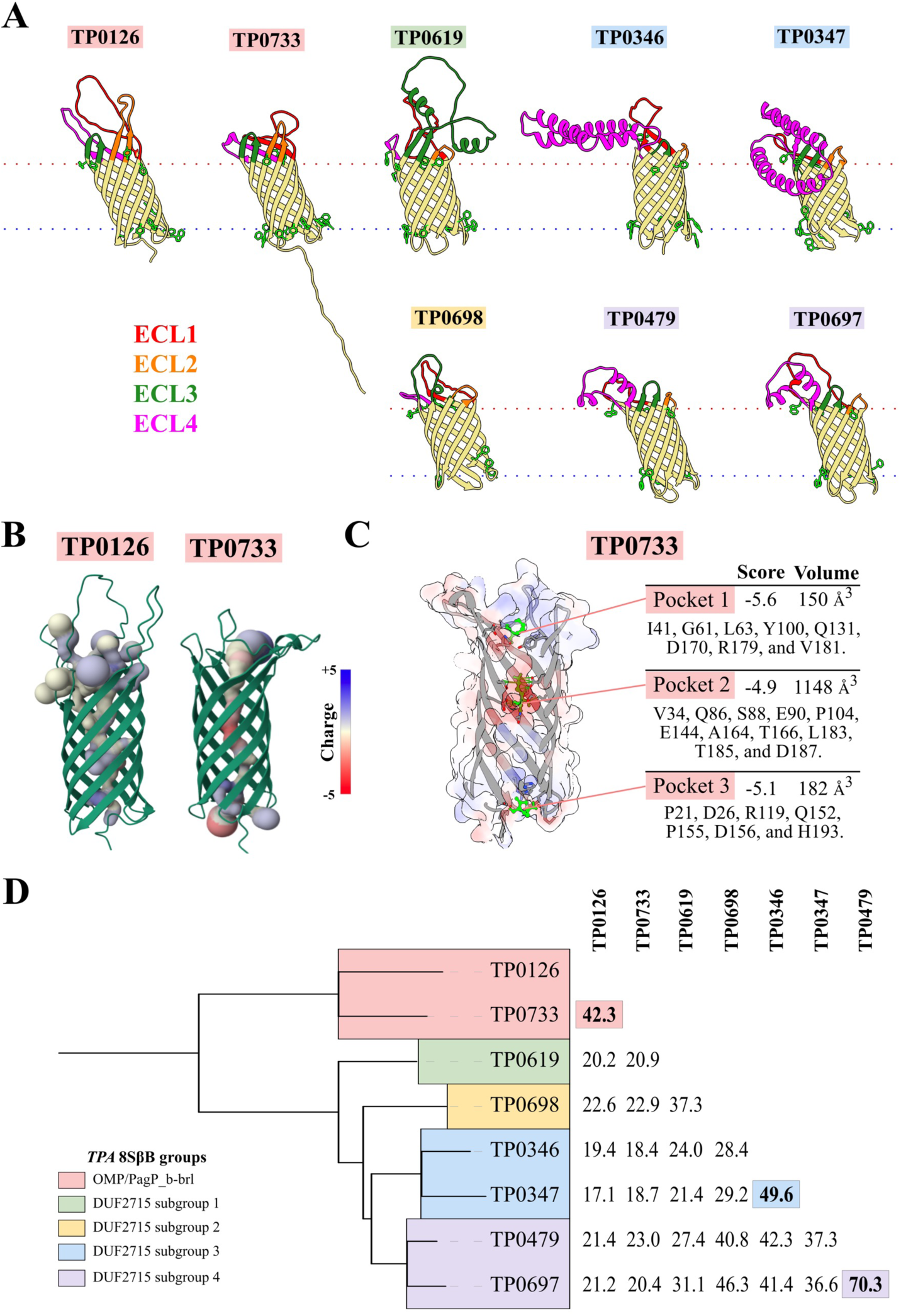
*TPA* 8SβBs segregate into subgroups with conserved β-barrel topology. (A) High-confidence AlphaFold 3 structural models for *TPA* Nichols 8SβB proteins (lacking signal peptides) maintain a conserved β-barrel topology with a tilted orientation within the outer membrane guided by interactions of aromatic residues (bright green side chains) at the bilayer interface. ECL1, red; ECL2, orange; ECL3, dark green; and ECL4, magenta. (B) Based on predictions by MOLEonline [29], OMP/PagP_b-brl, but not DUF2715-domain orthologs, form channels with different electrostatic properties that extend the lengths of the respective barrels. (C) Docking analysis using CB-Dock3 predicted three potential binding pockets for L-ornithine (CID: 6262) within TP0733. All pockets contained several polar and charged residues with predicted Vina scores ranging from −4.9 to −5.6 kcal/mol. (D) OMP/PagP_b-brl proteins TP0126 and TP0733 share only 42.3% protein similarity, while the DUF2715-domain proteins display wide variability (14.8 - 70.3%), forming four subgroups.

Although the β-barrel domains for *TPA* 8SβBs are highly similar, their predicted ECLs exhibit substantial structural diversification (Fig 1A). ECLs for the OMP/PagP_b-brl orthologs TP0126 and TP0733 are predicted to be disordered with wide variation in length. Except for TP0698, the DUF2715 orthologs each contain at least one α-helix-rich ECL that differs in size (30 - 85 aa) and position between orthologs. Phylogenetic comparison of the eight mature proteins revealed low to moderate overall amino acid similarity (17-70%), with the OMP/PagP_b-brl and DUF2715 proteins forming distinct branches. The six DUF2715 orthologs further separated into four subgroups, with higher structural similarity observed within subgroups 3 and 4 (Fig 1D).

Predicted structures for *TPA* 8SβBs also differ markedly from the solved structures of well-characterized prototypical 8SβBs (S5 Fig). While two *TPA* 8SβBs (TP0126 and TP0733) contain PagP domains, named after the lipid A palmitoyltransferase PagP, they lack distinguishing features of its namesake - the hydrophobic pocket required for interaction with palmitate, the conserved amphipathic residues (His, Asp, Ser) important for the catalytic activity, and the α-helix required for stabilization in the membrane [16, 32]. In contrast to the hydrophobic lumens of the substrate transporters OmpW [12] and CarO [31], the interiors for *TPA* 8SβBs are hydrophilic (S4 Fig). The α-helix-rich topology of *TPA* 8SβB ECLs also differs strikingly from the surface-exposed, extended β-strands for 8SβBs that mediate substrate binding (*e.g*., *Pseudomonas aeruginosa* CarO [13]) and interactions with host molecules (*e.g*., *Yersinia pestis* Ail [33]). The absence of a periplasmic domain for all but one (TP0733) *TPA* 8SβB ortholog further distinguishes them from bipartite 8SβBs (*e.g.*, *E. coli* OmpA [23]).

### Predicted interactions with host proteins implicate 8SβBs in adhesion to CEACAMs and extracellular matrix and complement evasion

As noted earlier, 8SβBs in bacterial pathogens serve multiple virulence-related functions [11]. Interactions have been reported between *Neisseria* spp. Opa proteins and CEACAM1, CEACAM3, and CEACAM5 [17]. Similarly, *Yersinia* spp. Ail had well-characterized interactions with the hemopexin domain of vitronectin [18], and domains 6-7 of complement inhibitor factor H [19]. To determine whether *TPA* 8SβBs exhibit similar host-binding activities, we next used AF3 to generate structural complexes between *TPA* 8SβBs and host molecules previously reported to interact with *N. gonorrhoeae* Opa (CEACAMs) and *Y. pestis* Ail (vitronectin and factor H). The biological likelihood and stability of the modeled protein-protein interfaces were assessed using PIZSA [34]; complexes with *Z*-scores >1.8 were analyzed with PRODIGY to estimate binding free energies (ΔG) and dissociation constants (*K_d_*) [35]. To validate our approach, we modeled the interactions of *N. gonorrhoeae* Opa60 and *Y. pestis* Ail (positive controls) and the β-barrel of *E. coli* OmpA (negative control). As expected, Opa and Ail generated high-confidence interfaces with their known ligands (CEACAMs 1, 5 and 6 for Opa60 and vitronectin and factor H for Ail), while OmpA failed to generate high-confidence complexes with any of the targets (S6A Fig). Although AF3 predicted complexes for all *TPA* 8SβBs except TP0733, only a subset displayed *Z*-scores above the cutoff (S6A Fig). In all these complexes, multiple *TPA* 8SβB ECLs contributed to the predicted binding interface, resulting in high-affinity interactions. DUF2715 subgroup 3 (TP0346 and TP0347) and subgroup 4 (TP0479 and TP0697) proteins displayed protein-CEACAM interfaces comparable to those for Opa60. TP0479 exhibited high affinities (ΔG = −9.6 to −10.4 kcal/mol) for all three CEACAMs tested, whereas TP0346, TP0347, and TP0697 preferentially recognized CEACAM1 (ΔG = −9 kcal/mol), CEACAM6 (ΔG = −8.4 kcal/mol), and CEACAM5 (ΔG = −9.7 kcal/mol), respectively (Fig 2 and S6B Fig). Five *TPA* 8SβBs displayed high-confidence interactions with extracellular matrix and complement-associated proteins. TP0619 exhibited the highest affinity overall, interacting with vitronectin (ΔG = −20.6 kcal/mol), whereas TP0698 interacted exclusively with the complement inhibitor factor H (ΔG = −11.9 kcal/mol) (Fig 2). TP0126 generated high-confidence interfaces with both vitronectin and factor H, but with lower predicted affinities than TP0698.

**Fig 2.**
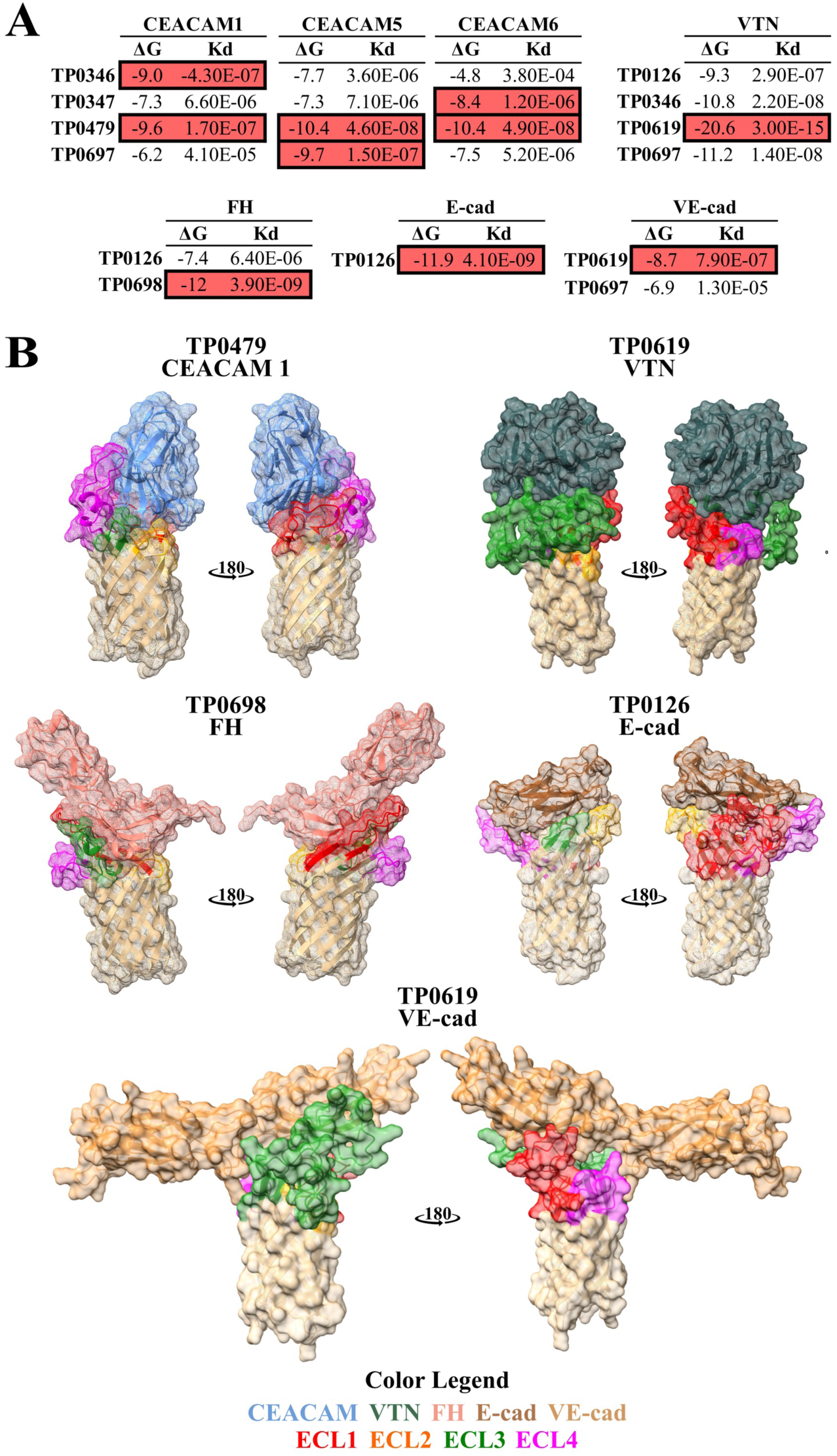
AlphaFold 3 predicts high-confidence interactions between *TPA* 8SβB and host proteins involved in attachment and complement evasion. (A) Predicted binding free energies (ΔG, kcal/mol) and dissociation constants (Kd, M) estimated by PRODIGY for high-confidence complexes between *TPA* 8SβB and human CEACAM1, CEACAM5, CEACAM6, the ninth fibronectin type III repeat (FNDC9), the vitronectin hemopexin domain (VTN), complement factor H sushi domains 6-7 (FH), E-cadherin (PDB: 1O6S), or VE-cadherin domains 4 and 5. The strongest predicted interaction for each host protein is highlighted in red. (B) Structural models of the highest-confidence complexes.

Interactions with cadherins within the epithelium (E-cadherin) and vascular endothelium (VE-cadherin) have also been implicated in pathogen attachment, invasion, and dissemination. Given the paucity of other surface-exposed proteins in *TPA*, we next hypothesized that 8SβBs may also contribute to cadherin-mediated host-cell interactions [36]. We therefore extended our analyses to explore *TPA* 8SβBs interactions with E-cadherin and VE-cadherin. In fact, TP0126 was predicted to interact with a high affinity to E-cadherin (ΔG = −11.9 kcal/mol), whereas TP0619 and TP0697 were predicted to interact with VE-cadherin, with TP0619 displaying the higher predicted affinity (ΔG = −8.7 kcal/mol) (Fig 2 and S6B Fig).

### Eight-stranded β-barrels are transcribed at markedly different levels during syphilitic infection of rabbits

As a starting point for elucidating how *TPA* orchestrates expression of its 8SβBs in the service of its parasitic strategy, we used hybridization capture RNA-seq [37] to determine their *in vivo* transcriptional profiles in spirochetes harvested at peak orchitis from rabbits infected intratesticularly with *TPA* Nichols or SS14. *tp0326/bamA*, encoding the β-barrel assembly machinery A (BamA) ortholog [38], and *tp0574/tpn47*, encoding the highly expressed 47-kDa lipoprotein [39], served as comparators. Transcripts for all eight 8SβBs were detected in both reference strains at levels comparable to or higher than *bamA* but substantially lower than *tpn47* (Fig 3). Overall, we saw a broad range of transcription, with *tp0697* and *tp0698* being the most highly expressed in both reference strains. In general, transcripts for the OMP/PagP_b-brl genes *tp0126* and *tp0733* were detected at lower levels than their DUF2715 counterparts. Four DUF2715 genes (*tp0346*, *tp0347*, *tp0697* and *tp0698)* were expressed at significantly higher levels (1.31-to 2.56-fold; *p* < 0.05) in *TPA* Nichols compared to SS14 (Fig 3). Visualization of mapped reads suggested that *tp0126*, *tp0127\**, *tp0315\**, *tp0346*, *tp0347* and *tp0733* are monocistronic, while the expression profiles of *tp0619*-*tp0618\** and *tp0698-tp0697* are less clear (Fig 4). *tp0479* appears to be transcribed as the first gene in an operon that includes *tp0478/zwf*, encoding the glycolytic enzyme glucose-6-phosphate dehydrogenase, and *tp0477/pgl*, encoding the pentose phosphate pathway enzyme 6-phosphogluconolactonase. Notably, *tp0127\**, *tp0315\** and *tp0619/tp0618\** reside in proximity to genes encoding members of *Treponema* repeat proteins (Tpr) subfamilies I (*tprD*, *tprF* and *tprI*) and II (*tprE*, *tprG* and *tprJ*) [40, 41] (Fig 4). Although *tp0315\** and *tp0619/tp0618\** are located immediately downstream and in the same orientation as *tprG/tprF* and *tprJ/tprI*, respectively, members of these two paralogous families do not appear to be co-transcribed.

**Fig 3.**
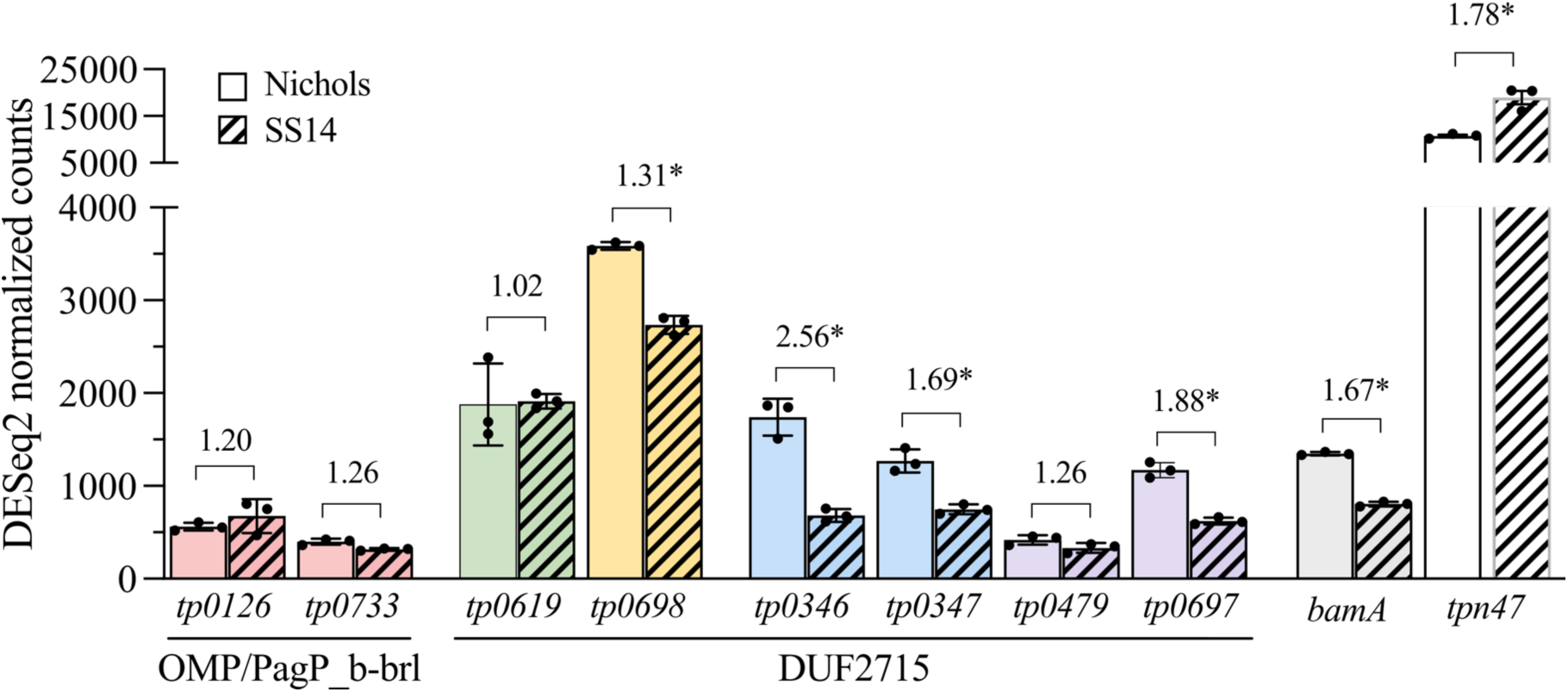
RNA-seq reveals substantial differences in transcription levels of *TPA* 8SβBs during syphilitic infection of rabbits. Bars represent mean DESeq2-normalized read counts (± SD) for 8SβB genes in *TPA* Nichols (solid bars) and SS14 (hatched bars) harvested from rabbit testes at peak orchitis. Dots indicate biological replicates (n = 3 per strain). Transcript values for *bamA* (*tp0326*) and *tpn47* (*tp0574*) are shown as comparators. Numbers above bars indicate DESeq2-estimated fold changes in transcript abundance between strains. Asterisks (*) denote genes exhibiting significant (*q* < 0.05) differences in expression between strains.

**Fig 4.**
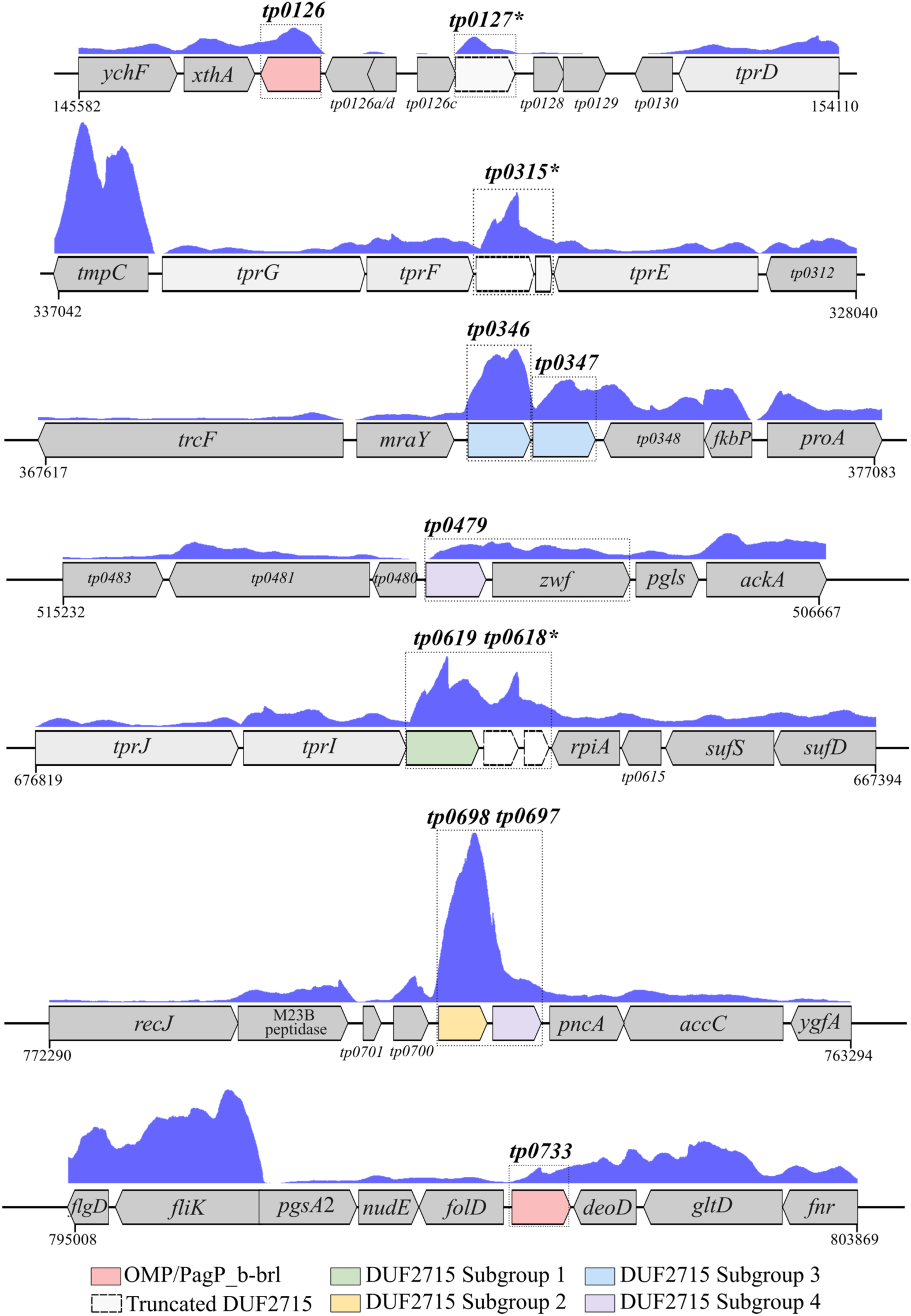
RNA-seq mapped read coverage suggests that most *TPA* 8SβB loci are monocistronic. Coverage tracks (blue) represent total RNA-seq reads mapped to the corresponding genomic regions from *TPA* Nichols harvested from rabbit testes. The *y*-axis scale for all coverage tracks was adjusted to a maximum of 7,000 read counts. Dashed boxes indicate predicted transcribed regions for the corresponding 8SβB-encoding gene. Detailed descriptions of all genes are provided in S1 Table.

### *TPA* 8SβBs ECLs differ markedly in immunogenicity during rabbit infection

Rabbits infected with *TPA* develop long-lasting immunity to reinfection [42, 43], with protection widely thought to be mediated by antibodies directed against the spirochete’s rare OMPs [42, 44–46]. To determine whether *TPA* 8SβBs elicit antibody responses that could potentially contribute to infection-induced protective immunity, we focused on their antibody accessible ECLs. ElliPro [47] predicted that all eight 8SβBs contain one or more ECLs harboring linear and/or discontinuous B-cell epitopes (BCEs) (Fig 5 and S2 Table). To assess whether these predictions reflect antibody production during infection, we examined the reactivity of IRS with individual ECLs displayed in a native-like conformation by *Pyrococcus furiosus* thioredoxin (*Pf*Trx) scaffolds [48, 49]. Heterogeneity in ECL-specific antibody responses between immune rabbits was evident. As shown in Fig 5, little (AUC < 2.0) to no reactivity was observed for ECLs from TP0126, TP0346, TP0347 and TP0619, despite the presence of multiple high-confidence BCEs. We saw the strongest reactivity against TP0698 (ECLs 1, 3, and 4), the most highly expressed ortholog, and TP0733 (ECL1), one of the lowest expressed. Surprisingly, given that 8SβB-encoding genes were expressed at comparable or higher levels in *TPA* Nichols compared to SS14 (Fig 3), ECL-specific responses trended higher in rabbits infected with the latter strain (Fig 5).

**Fig 5.**
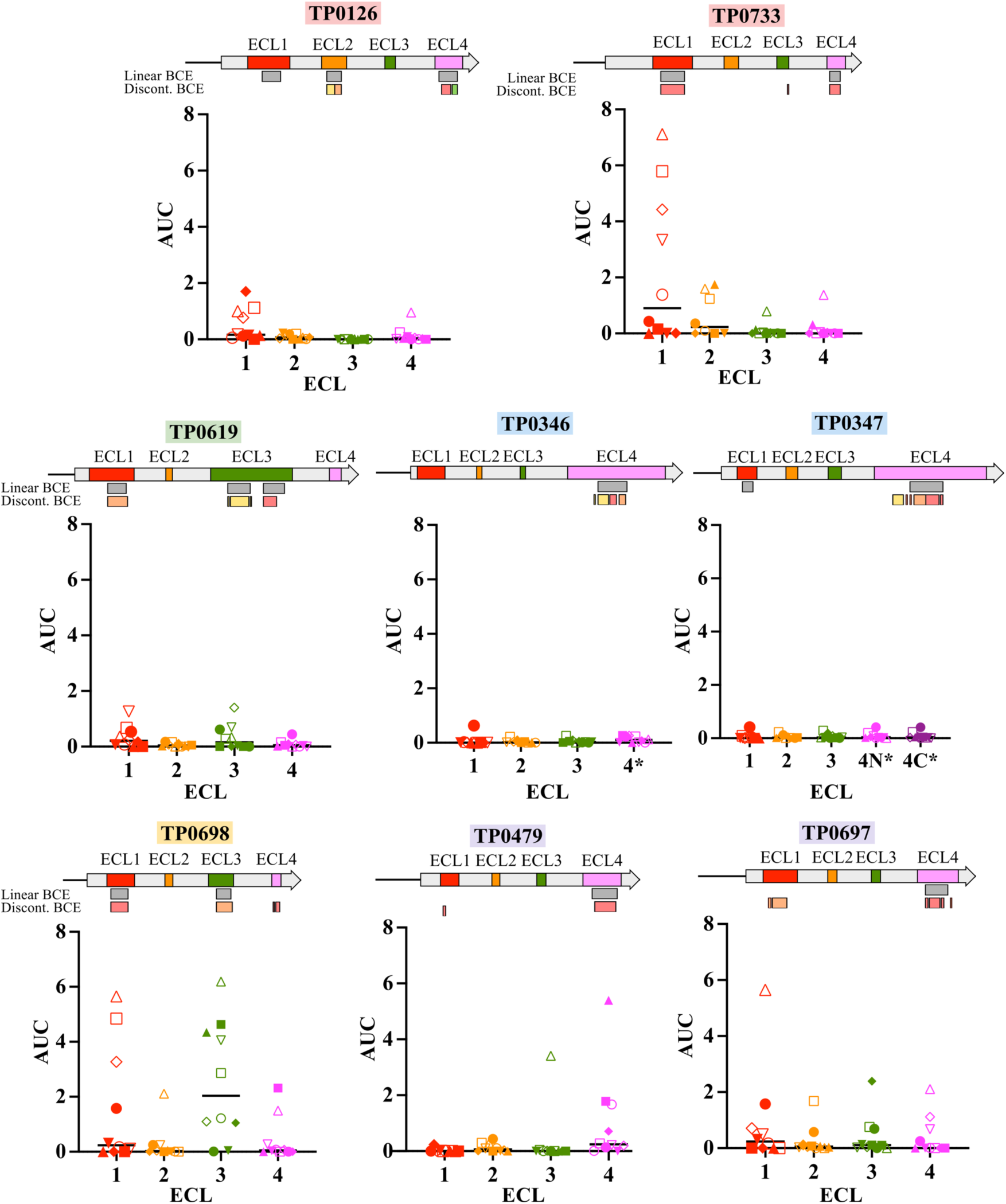
*TPA* 8SβBs show a broad spectrum of ECL antigenicity profiles with immune rabbit sera. Reactivity by ELISA (right) of *Pf*Trx-scaffolded 8SβB ECLs against sera from *TPA* Nichols (filled symbols) and SS14 (open symbols) immune rabbits; *n* = 5 animals per strain; each animal is represented by a different symbol. ELISA reactivity was measured as AUC (see Materials and Methods). ECL1, red; ECL2, orange; ECL3, green; and ECL4, magenta. Linear (grey) and discontinuous (red, orange, yellow and green) B-cell epitopes (BCEs) were predicted by ElliPro using a cutoff of 0.8 (Supplementary Table 2). Regions from the same discontinuous BCE are similarly colored. Due to the large size of ECL4 in TP0346 (70 aa) and TP0347 (82 aa), single helical regions containing predicted BCEs were displayed on independent *Pf*Trx constructs and designated 4* for TP0346 ECL4 and 4N* and 4C* for the N-terminal and C-terminal helices of TP0347 ECL4, respectively.

### 8SβBs are highly conserved across geographically diverse strains and subject to purifying selection

To better understand how 8SβBs contribute to persistence of *TPA* within endemic populations, we examined sequence variability of the Nichols orthologs across 1706 *TPA* genomes (483 Nichols-lineage [28%] and 1,223 SS14-lineage [72%]) obtained from patients in Africa, Asia, Europe, Oceania, North and South America (S3 Table). Overall, we saw very low variability among the eight 8SβBs when compared to the *TPA* Nichols reference strain. TP0698, the most highly transcribed (Fig 3) and immunogenic 8SβB (Fig 5), was invariant. Forty amino acid substitutions were identified across all the remaining 8SβB loci (S4 Table). Four substitutions (two in ECLs 1 and 4 of TP0346 and two in β-strand 4 of TP0619) segregated almost exclusively with clinical strains belonging to the SS14-lineage, and therefore, were found at higher frequency (∼72% of genomes) (Fig 6A and S4 Table). Of the remaining 36 substitutions, only four (one in ECL4 of TP0346, two in ECL1 and one in ECL4 of TP0733) were found in >1% of the genomes (Fig 6A and S4 Table). All four high-prevalence substitutions mapped to ECLs containing predicted BCEs (Fig 5). Notably, almost all clinical strains encoding these substitutions belonged to the Nichols-lineage (Fig 6B). We also noted geographic skewing for variability in TP0733 and TP0346; mutations in TP0733 ECL1 (N73D and G79S) were exclusive to strains from Africa, with 29% of strains from this region harboring either or both substitutions. The remaining mutations were distributed across multiple geographic regions, with the greatest prevalence in strains from Europe, where ∼14% carried substitutions in TP0346 ECL4 (K163E) and/or TP0733 ECL4 (S199P).

**Fig 6.**
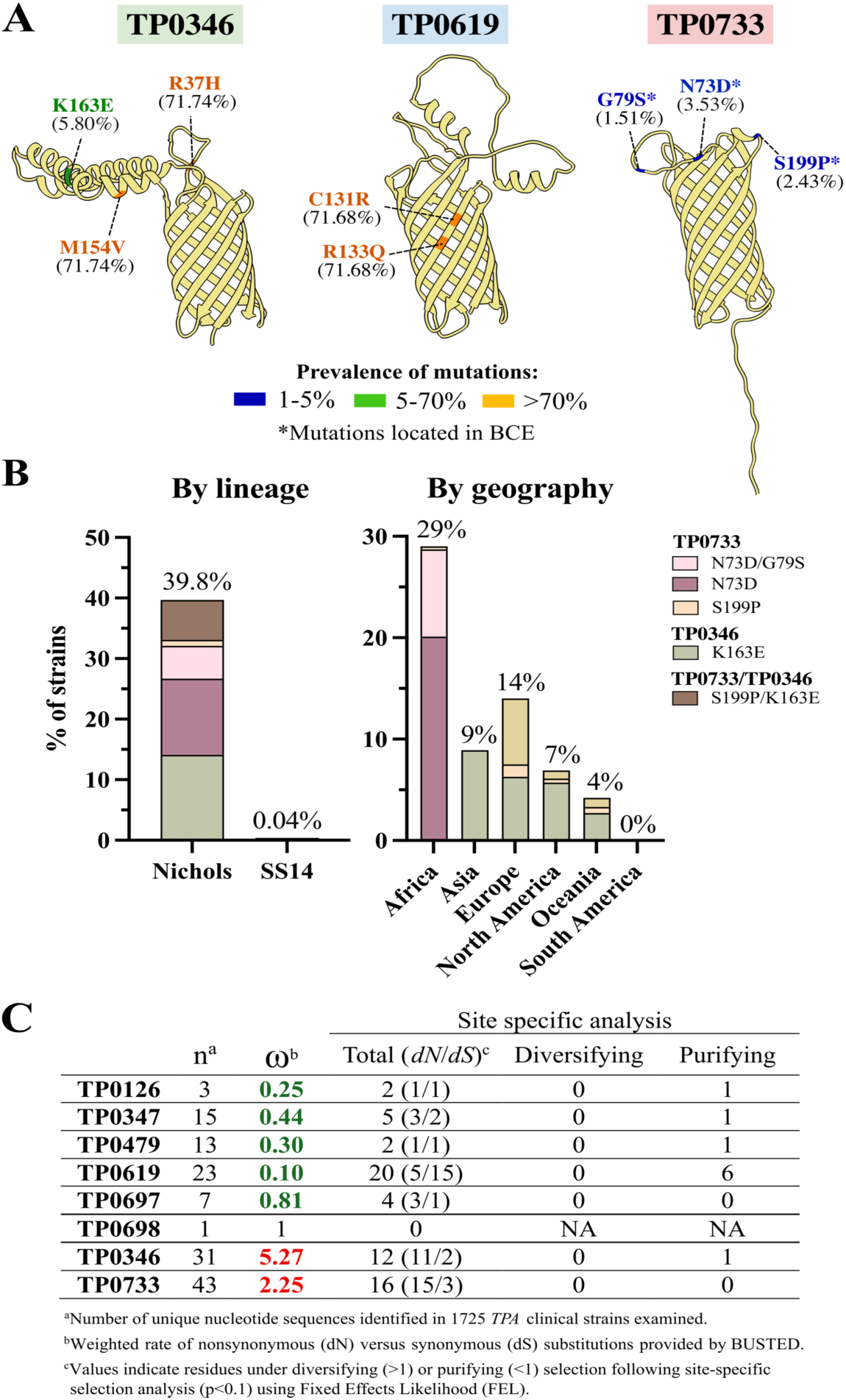
*TPA* 8SβBs are highly conserved across clinical strains from patients with early syphilis. (A) Among 1,706 analyzed clinical strains, only three 8SβB proteins contain residues that varied in more than 1% of strains. Only a single mutation (TP0346 K163E) was present in > 5% of clinical strains (green). Variants present in >71% of clinical strains (orange) represent lineage-specific differences. (B) Mutations were highly influenced by lineage and geography location of samples. (C) Residues under selective pressure were identified using maximum likelihood-based analyses available on the Datamonkey webserver. Gene-wide selection was assessed using BUSTED (Branch-site Unrestricted Statistical Test for Episodic Diversification), while residues under diversifying selection were identified using FEL (Fixed Effects Likelihood) with a *p*-value threshold of 0.1. NA, not applicable. Detailed information about codon variation in individual strains is available in S5 Table.

The low frequency of non-synonymous mutations in *TPA* 8SβBs suggested that the corresponding genes are subject to purifying selection. To examine this possibility, we used maximum likelihood-based models to estimate the ratio (ω) of non-synonymous (dN) to synonymous (dS) nucleotide substitutions within 8SβB-encoding genes. Five (*tp0126*, *tp0347*, *tp0479*, *tp0619* and *tp0697*) contained ω values ranging between 0.1 and 0.8, consistent with gene-wide purifying selection (Fig 6C). Site-specific analyses identified ten sites (one each in TP0126, TP0346, TP0347, and TP0479 and six in TP0619) under significant (*p* < 0.01) purifying selection. Although positive ω values were observed for the two most variable 8SβBs, TP0346 (ω = 5.27) and TP0733 (ω = 2.25), neither contained nucleotide residues under significant diversifying selection. TP0698 was invariant at the nucleotide as well as the amino acid level.

### Hypermutable polyG tracts expand the 8SβB repertoires in *TPA* clinical strains

Giacani *et al*. [50] previously reported that transcription of *tp0126* is regulated by phase variation due to slipped strand mispairing in a variable upstream polyguanine (polyG) tract. A survey of the *TPA* clinical genomes, described above, revealed that homopolymeric tracts (≥5 Gs) are widely distributed across 8SβB loci, occurring not only upstream but also within coding sequences (S7 Fig). PolyG tracts were identified upstream of genes encoding five full-length (TP0479, TP0347, TP0619, TP0697 and TP0698) and two truncated (*tp0127*\* and *tp0315*\*) 8SβBs. While the polyG tracts upstream of *tp0619*, *tp0127*\* and *tp0315*\* were invariant and short (5 bp), those upstream of the *tp0126, tp0347, tp0479* and *tp0697* coding sequences varied in length (7-16 Gs) between the Nichols and SS14 reference genomes and the clinical strains (Fig 7A and S8A Fig). The positions of the polyG tracts varied widely relative to the predicted ribosome-binding sites (23-46 bp) and translational start codons (10-53 bp) (Fig 7A).

**Fig 7.**
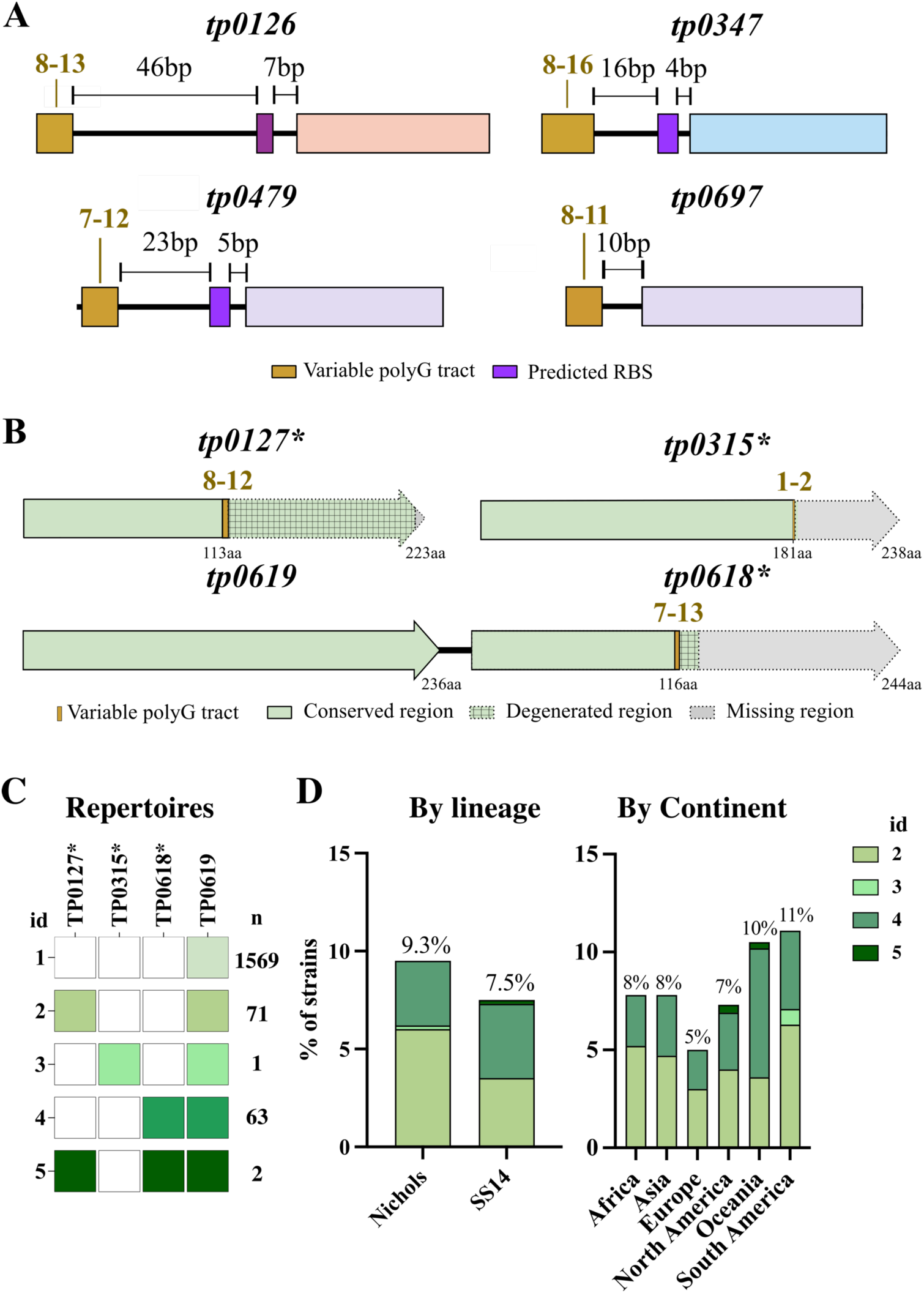
Variability in intragenic polyG tracts gives rise to different DUF2715 subgroup 1 repertoires. (A) Alignment of 8SβB coding regions from Nichols and SS14 reference strains and 1,706 clinical strains revealed 10 conserved (yellow lines) and 3 variable (red lines) polyG tracts within DUF2715 subgroup 1 orthologs; the numbers above each line indicate the number of guanosine residues within the corresponding tract. (B) Variations in polyG tracts generate five repertoires of full-length 8SβBs within clinical strains; repertoires in 137 strains differ from the reference strains. (C) Existence of variant 8SβB repertoires (2-5) appears not to be influenced by lineage or geographic location of strains. Detailed information about individual strains is available in S6 Table.

We also identified 21 intragenic polyG tracts within nine of the eleven 8SβB-encoding genes. While most were invariant, the polyG tracts within *tp0127*\*, *tp0315*\*, and *tp0618*\* varied (1-13 Gs) among clinical strains (Fig 7B and S8B Fig). In 137 clinical strains, insertions/deletions within the polyG tracts restored the open reading frames for *tp0127\** (71 strains), *tp0618\** (63 strains), *tp0315\** (1 strain), or both *tp0127\** and *tp0618\** (2 strains) (Fig 7C). The restored proteins displayed strong sequence and structural similarity with the DUF2715 subgroup I ortholog TP0619 (S9A and B Figs, respectively). Indeed, the restored TP0315* was identical to TP0619. Variability within subgroup 1 was largely confined to the ECLs, where substitutions and in-frame insertions frequently altered the electrostatic properties of surface-exposed regions (S9C Fig). Unlike TP0619/TP0315*, the restored TP0127* and TP0618* did not generate vitronectin complexes with the same confidence or predicted affinity as their subgroup partners TP0619 and TP0315*, suggesting further specialization within the subgroup (S9D Fig).

## Discussion

As an obligate human pathogen and extreme auxotroph, *TPA* must meet the diverse physiological and pathological demands of infection while minimizing immune recognition by the host to disseminate and persist [10]. The spirochete’s ability to balance these competing demands is facilitated in part by its remarkably slow replication rate (∼30 h) [51, 52], which reduces the need for high copy numbers of individual OMPs and contributes to its stealth pathogenicity. *TPA* differs from most Gram-negative bacteria in containing multiple paralogous OMP families [7, 9]. Within each family, individual proteins can diverge and specialize in function, expanding the functional capacity of the OMPeome [7]. The 8SβB family appears to be central to this survival strategy. The functional versatility of 8SβBs in other bacteria [11] suggests that diversification of this family in *TPA* could support multiple physiologic and virulence-related functions required for infection and persistence in humans. Our identification of four additional full-length 8SβBs substantially expanded the *TPA* OMPeome (Fig 8A), underscoring the presumptive contribution of these proteins to the spirochete’s pathogenic lifestyle (Fig 8B). Divergence within this structurally minimalist family occurs in both the β-barrels and ECLs, with predicted implications for channel formation and interaction with diverse host molecules, respectively. Notably, *TPA* 8SβBs have diverged extensively in both sequence and structure from Gram-negative orthologs characterized to date, as evidenced by their unique ECL organization and the emergence of the treponeme-specific DUF2715 domain. Despite an extraordinary degree of sequence conservation among clinical strains, the repertoire of 8SβBs in *TPA* appears to be expanding and diversifying *via* an unusual mechanism - variation in homopolymeric tracts - potentially contributing to the ability of syphilis spirochetes to thrive on a global scale.

**Fig 8.**
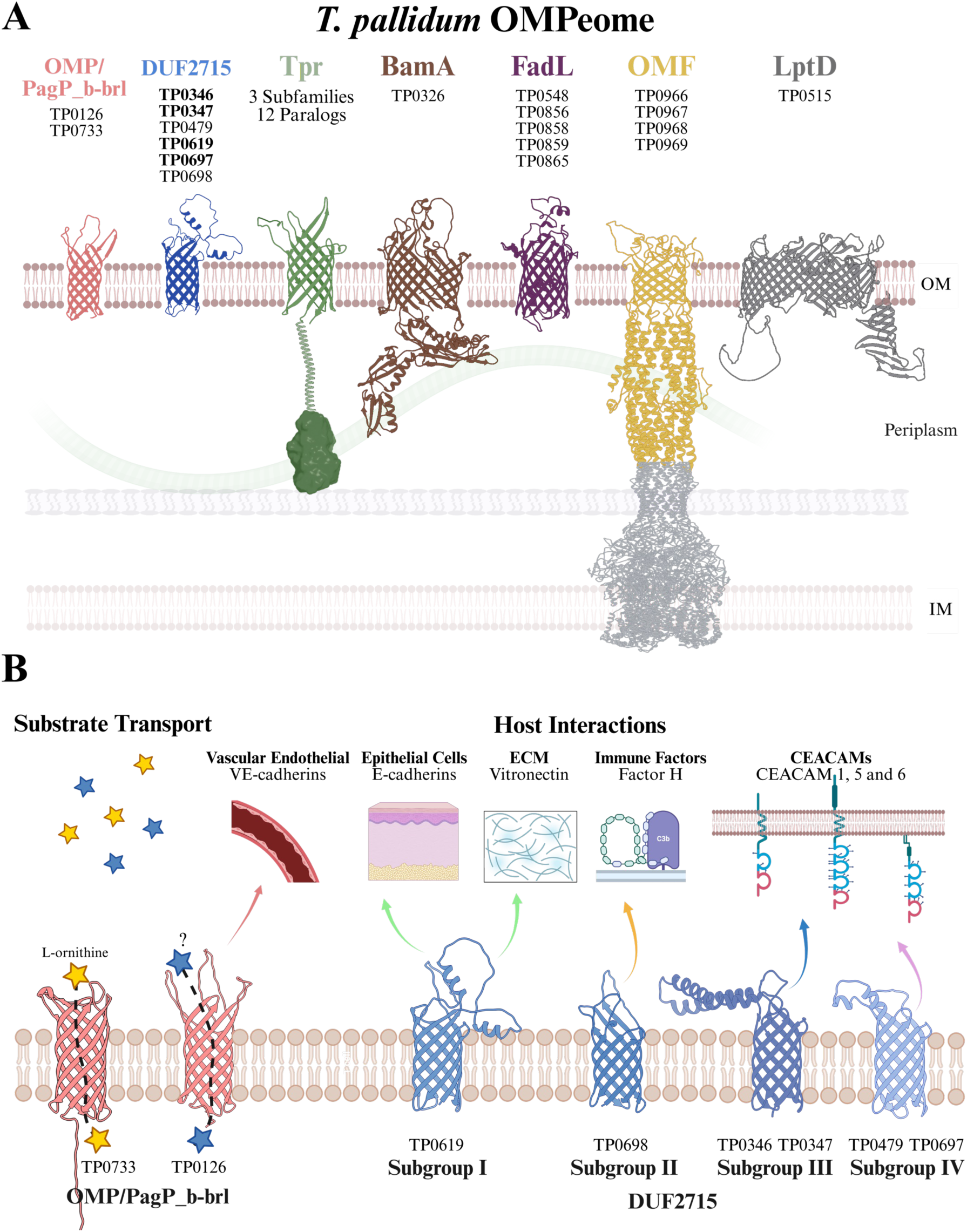
The expanded *TPA* OMPeome contains 31 OMPs. (A) The identification of four additional DUF2715-containing orthologs expands the predicted *TPA* OMPeome to 31 OMPs distributed across seven protein families (OMP/PagP_b-brl, DUF2715, Tpr, BamA, FadL, OMF, and LptD). Structural models were generated by Alphafold 3. (B) The functional versatile *TPA* 8SβB family may support multiple physiologic and virulence-related functions required for infection and persistence. Proposed functions remain to be experimentally confirmed.

Most substrate import in Gram-negative bacteria is mediated by trimeric 16-stranded porins and TonB-dependent receptors [53]. *TPA* lacks these OMP families [7, 9], indicating that other OMPs fulfill the essential nutrient-acquisition functions required to sustain its auxotrophic lifestyle. This divergence from the Gram-negative paradigm is exemplified by the porin activity reported for Tprs, a paralogous family of bipartite OMPs found exclusively in treponemes [40, 54]. Here, we propose that the OMP/PagP_b-brl 8SβBs (TP0126 and TP0733) contain predicted transmembrane channels and may likewise contribute to nutrient acquisition in *TPA*. Homology between these proteins and OmpW has previously been proposed [7, 27, 50]. However, the hydrophilic nature of the channels for these treponemal 8SβBs suggests that, unlike OmpWs, they may transport hydrophilic rather than hydrophobic substrates [12]. The distinct electrostatic properties between channels in TP0126 and TP0733 further support the transport of substrates with different charges. In *A. baumannii* CarO, charged residues selectively promote the uptake of ornithine and other positively charged substrates [31]. *TPA* requires ornithine as a structural component of its PG [30, 31] but lacks the enzymatic pathways for its *de novo* synthesis from arginine or glutamate [55, 56]. The abundance of charged residues throughout the TP0733 channel was predicted to favor interactions with ornithine, supporting a potential function analogous to *A. baumannii* CarO [31]. Notably, the small diameter of 8SβBs usually precludes formation of a constitutively open channel, requiring rearrangement of side chains within the barrel interior for substrate passage [57]. Therefore, the absence of predicted channels in DUF2715 orthologs does not exclude transport activity. In addition to its porin activity, OmpA contains a periplasmic domain that interacts with PG, tethering the OM and preserving cell envelope integrity [58]. TP0733 was the only *TPA* 8SβB containing a predicted periplasmic region, but, unlike the structured PG-binding domain of OmpA, this N-terminal extension was predicted to be disordered.

Several functionally characterized 8SβBs participate in host interactions, with the structure and composition of their ECLs determining their binding specificities for host molecules and receptors [11]. As syphilitic infection progresses, *TPA* must interact initially with epithelial surfaces and extracellular matrix at the portals of entry and then vascular endothelium and peripheral tissues during and after dissemination [3, 59–63]. Our structural predictions suggest that *TPA* 8SβBs contribute to these interactions and that differences in their ECLs determine specificity for distinct host molecules. Unlike extended β-strand topology observed in prototypical Gram-negative orthologs, most *TPA* 8SβB displayed one or more ECLs with an α-helix-rich topology, potentially creating interaction platforms that differ substantially from those of other 8SβBs [18, 64, 65]. The broad distribution of CEACAMs across epithelial, endothelial, and immune cells makes them attractive targets for colonization as well as invasion, with interactions promoting attachment and influencing tissue tropisms [66, 67]. In *N. gonorrhoeae*, the differential recognition of CEACAM1 and CEACAM5 by Opa proteins can influence the outcome of infection, promoting, respectively, long-term colonization of the lower genital tract and penetration into upper tract accompanied by acute inflammation [68]. *TPA* orthologs from DUF2715 subgroups III (TP0346 and TP0347) and IV (TP0479 and TP0697) were predicted to interact with CEACAMs. Notably, each was predicted to bind with higher affinity to a different CEACAM, suggesting that individual *TPA* 8SβBs may be preferentially recruited at distinct sites or stages of infection. Interactions of *N. gonorrhoeae* Opa with the transmembrane signaling CEACAM1 expressed on epithelial cells [69], as well as on T-[70] and B-[71] lymphocytes, modulate innate and adaptive immune responses and contribute to bacterial persistence [17]. Such interactions could be particularly relevant for *TPA*, which disseminates rapidly to lymphoid tissues, where it can persist and disrupt immune responses [3, 72]. Several pathogens also exploit interactions mediated by 8SβBs to inhibit complement activation, as exemplified by those of *Y. pestis* Ail [18] and *Rickettsia conorii* Adr1 [73] with the multifunctional protein vitronectin, and of *Salmonella* spp. Rck [20] with complement inhibitor factor H. *TPA* DUF2715 subgroups I (TP0619) and II (TP0698) 8SβBs may similarly engage with these molecules and interfere with complement activation. As essential components of cell junctions [74], cadherins also represent plausible receptors for *TPA* during hematogenous dissemination. Notably, *TPA* is one of few pathogens able to cross the placenta [75]. In *Listeria monocytogenes,* the internalin InlA interacts with E-cadherins expressed by syncytiotrophoblasts to promote invasion of the placental barrier [76]. Likewise, *TPA* can disrupt VE-cadherin intercellular junctions as a mechanism to traverse endothelial barriers [77]. Although interactions between 8SβBs and cadherins have not previously been reported in other bacteria, the paucity of surface-exposed proteins in *TPA* raises the possibility that 8SβBs have evolved to perform this function. Our structural predictions support specific OMP interactions that can be experimentally investigated to address the long-standing question of the molecular mechanisms underlying *TPA-*host interactions.

As noted above, *TPA* 8SβBs appear to have evolved specialized physiological and virulence-related functions, potentially requiring the expression of individual OMPs to be calibrated within specific host milieus. Consistent with this notion, we observed that *TPA* 8SβB genes are transcribed at markedly different levels in rabbit testes. Notably, transcript levels for all eight were consistently higher in the Nichols strain compared to SS14. Although the basis for these differences remains unclear, a similar pattern was observed for the essential OMP assembly factor BamA, suggesting that increased OMP transcription may reflect the higher replication rate of the Nichols strain [78]. In addition to intrinsic differences in promoter activity, variability within upstream homopolymeric tracts may influence transcription and, consequently, host adaptation [79]. In *N. meningitidis*, variation in a polyG tract located between the −10 and −35 promoter elements for *porA* is proposed to alter RNA polymerase (RNAP) binding, affecting gene expression and contributing to immune evasion [80]. In *TPA*, polyG tracts have been implicated in transcriptional regulation of *tp0126* [50] and *tpr* [81] genes. The polyG tract upstream of *tp0126* lies within its predicted RNAP binding region and presumably acts similarly to that described for *porA* [50], while the polyG tracts upstream of the *tprs* are much closer to their transcriptional start sites (TSS), potentially altering RNAP binding and/or the initiation of transcription, with shorter polyG tracts enhancing expression [81]. However, Giacani *et al*. [81] observed only a partial correlation between polyG length and mRNA levels for *tpr*s in rabbit testes. Herein, we report previously unrecognized polyG tracts upstream of several DUF2715-encoding genes; unlike *tp0126*, these are positioned near the predicted TSSs for DUG2715 genes and may act through a mechanism similar to that proposed for *tpr*s. The length of polyG tracts varied between Nichols and SS14 reference strains and across clinical strains belonging to either lineage. Notably, the polyG tract upstream of *tp0347* was considerably shorter (−3 nucleotides) in SS14-lineage strains, yet transcriptional levels were higher in Nichols, contrasting with the regulatory mechanism proposed by Giacani *et al*.

The well-recognized ability of *TPA* to disseminate and evade immune responses, attributed in part to the paucity of OMPs and lack of surface-exposed PAMPs, has earned the spirochete the designation ‘the stealth pathogen’ [10]. Nevertheless, studies dating to the pre-molecular era demonstrated that rabbits infected with *TPA* can develop complete protective immunity [42, 43], with functional antibodies (*i.e*., opsonic and/or neutralizing) believed to be correlates of protection [44–46]. Identifying the ECLs targeted by these protective antibodies should therefore provide valuable insights into the mechanisms underlying spirochetal persistence in humans and inform vaccine development. Using the *Pf*Trx scaffold [82] to display the individual ECLs for all eight *TPA* 8SβBs in a native-like conformation, we observed a striking heterogeneity in ECL-specific antibody responses in sera from immune rabbits. Only three ECLs (ECL1 of TP0733 and ECLs 1 and 3 of TP0698) exhibited strong antigenicity that could potentially contribute to protective immunity, whereas other ECLs failed to induce appreciable antibody responses. The basis for these differences is puzzling. Observed antigenicity does not appear to correlate with their predicted antigenic potential, as several ECLs from the poorly antigenic TP0126 and TP0619 contained predicted strong BCEs. High antigenicity was also not simply a consequence of transcriptional abundance; the two most antigenic orthologs, TP0698 and TP0733, were the most and least transcribed 8SβBs, respectively. Moreover, *TPA* 8SβBs were transcribed at lower levels in the SS14 strain, yet sera from rabbits infected with this strain generally exhibited higher reactivity than rabbits infected with the Nichols strain. Future immunization experiments with scaffolded ECLs will determine if these dichotomies reflect intrinsic differences in antigenicity, irrespective of BCE predictions, or how *TPA* strains engage the adaptive immune system during syphilitic infection. How closely human responses to these ECLs parallel those in rabbits also remains to be determined. The extraordinarily high sequence conservation of individual *TPA* 8SβBs orthologs within clinical strains suggests that, in general, they are not subject to a high degree of immune pressure either because they are not inducing antibodies or, as observed for TP0698, the antibodies that are induced are of insufficient quality to overcome functional constraints that would resist the introduction of non-conservative substitutions. Our finding that most members of this family are under purifying selection lends credence to the latter possibility.

Sequence variation in antibody-accessible regions, typically ECLs, is a fundamental mechanism for generating bacterial phenotypic diversity at the host-pathogen interface [83]. By promoting evasion of antibody-mediated responses and enhancing the host-interaction properties of OMPs, mutations can profoundly impact the survival and proliferation of pathogens within individuals and susceptible populations [84–87]. Recombination also serves as a major driver of OMP variability [88]. Most notably, recombination within the ECLs of *N. gonorrhoeae* Opa generates mosaic repertoires that can alter tissue tropisms and contribute to immune evasion [84]. Additionally, nonsynonymous single-nucleotide polymorphisms (SNPs) can have substantial functional consequences, as illustrated by the *in vivo*-selected point mutations in ECL4 of PorA that promoted novel tissue tropisms in *Campylobacter jejuni* [85]. In *TPA*, recombination has contributed substantially to diversification, with most reported events, such as those in BamA and the FadL TP0548, involving ancestral *TPA* lineages and endemic treponemes (*T. pallidium* subspecies *endemicum* [*TEN*] and *pertenue [TPE]*) [89, 90]. The extent to which recombination contributes to OMP diversification in contemporary *TPA* strains remains unclear, but there is evidence that such events continue to occur [89, 91–95]; recombination involving the FadL-like gene *tp0865* was recently described in a case report of a patient coinfected with two *TPA* strains [96]. SNPs, however, appear to be the predominant drivers of *TPA* OMP diversification within circulating *TPA* strains [90, 92, 97]. Our previous analysis of BamA and FadLs in geographically diverse *TPA* strains identified a high number of non-synonymous mutations occurring in the extracellular regions of these proteins. Notably, mutations were more prevalent within the Nichols lineage and several segregated according to the geographic origin of the strain [90]; of particular interest, individual proteins within the FadL family appear to be subject to different degrees of immune pressure. Herein we found that *TPA* 8SβBs exhibit, overall, substantially less sequence diversity than BamA and FadLs. The four prevalent mutations identified in TP0346 and TP0733, however, showed similar patterns of distribution to those found in other OMP families; mutations occurred predominantly within the Nichols lineage, while two (N73D and G79S) were found exclusively in strains from Africa.

As exemplified by studies in *N. gonorrhoeae* [98] and *Bordetella pertussis* [99], hypermutable polyG tracts can generate phenotypic heterogeneity by (i) reversibly switching open reading frames between ‘on’ and ‘off’ states within a population and (ii) generating stable sequence variants that are maintained across generations. While we cannot completely rule out a regulatory role for the upstream polyG tracts, our analysis of intragenic polyG tracts within *TPA* 8SβBs provides strong support for the latter mechanism. Of the 1706 *TPA* clinical strains examined, 137 contained insertions/deletions within subgroup I DUF2715 orthologs (TP0127 [71 strains], TP0315 [1 strain], TP0618 [63 strains] and both TP0127 and TP0618 [2 strains]) that restored the corresponding open reading frame. TP0619, the sole subgroup 1 member in *TPA* Nichols and SS14 reference strains, is predicted to interact preferentially with vitronectin and VE-cadherin. Thus, expanding subgroup 1 proteins may enhance, or even extend, the functionality of these 8SβBs. It is tempting to speculate that we have caught *TPA* in the act of expanding its OMP repertoire, capturing an evolutionary process that enables the spirochete’s ability to thrive within and spread across at-risk human populations. Longitudinal population-based genomic studies of *TPA* circulating globally should enable us to track the emergence of clinical strains with alternative OMP repertoires and determine whether this expansion confers a selective advantage. The development of systems for the continuous cultivation *in vitro* [100] and genetic manipulation [101] of *TPA* provide a biological framework to dissect the functions of individual OMPs and examine how OMPeome diversity contributes to competitive fitness, dissemination, and persistence of *TPA* during natural infection.

## Materials and Methods

### Ethics Statement

All animal experimentation was conducted in accordance with “The Guide for the Care and Use of Laboratory Animals (Eighth Edition)” [102] using protocols reviewed and approved by the UConn Health Institutional Animal Care and Use Committee (Animal Welfare Assurance [AWA] number A347-01).

### Intratesticular propagation of *TPA* in rabbits and generation of immune rabbit sera

*TPA* Nichols and SS14 reference strains were propagated by intratesticular inoculation of adult male New Zealand White (NZW) rabbits as previously described [103]. Treponemes were harvested at peak orchitis in 0.5-1 ml of CMRL medium (ThermoFisher Scientific, Inc., Waltham, MA, USA) supplemented with 10% heat-inactivated normal rabbit sera (NRS). After 2 h, treponemes were recovered and enumerated by darkfield microscopy using a Petroff-Hausser counting chamber (Hausser Scientific, Horsham, PA, USA). Immune rabbit sera (IRS) against *TPA* Nichols and SS14 reference strains were generated by inoculation of rapid plasma reagin (RPR)-nonreactive adult male NZW rabbits as previously described [49].

### Identification of 8-stranded β-barrel OMP orthologs in *Treponema pallidum* subspecies *pallidum*

Previously unrecognized 8SβB OMPs within the *TPA* Nichols (UP000000811) and SS14 (UP000001202) proteomes were identified by searching the membrane β-barrel (MBB) clan (CL0193) within the EMBL-EBI InterPro database [104]. Corresponding coding sequences were obtained from *TPA* Nichols (NC_021490.2) and SS14 (NC_021508) reference genomes in NCBI. Predicted signal peptides and signal peptidase I (SPaseI) cleavage sites within coding sequences (S1 Fig) were identified based on SignalP (ver. 6.0) [105]. Translational start sites for 8SβB orthologs were curated manually based on the position of predicted SPaseI cleavage sites, putative ribosome binding sites, and/or RNA-seq mapped read data (S2 Fig). Signal peptide sequences were removed before phylogenetic and structural analyses.

### Multiple sequence alignments and phylogenetic comparisons

Identity and similarity among 8SβB orthologs within *TPA* were based on multiple sequence alignments generated using MAFFT (ver. 7.525) [106]. Maximum likelihood phylogenies were calculated using PhyML (ver. 3.0) [107, 108]. Tree improvement was performed by subtree pruning and regrafting (SPR) method with ten random starting trees. Robustness of branches was assessed by an approximate likelihood-ratio test (aLRT) [109]. The resulting trees were visualized and annotated using Interactive Tree of Life (ver. 6.8.1) [110].

### Structure-based analyses

Conserved domain searches were performed using InterPro [104]. Three-dimensional structural models of *TPA* 8SβB proteins, with signal peptide sequences removed, were generated using AlphaFold 3 [111], as either individual proteins or in complex with human CEACAM1 (UniProt: P13688), CEACAM5 (UniProt: P06731), CEACAM6 (UniProt: P40199), the vitronectin hemopexin-like domain (PDB: 6O5E), complement factor H domains 6 and 7 (UniProt: P08603), E-cadherin (PDB: 1O6S), or VE-cadherin domains 4 and 5 (UniProt: P33151). Simulations of protein positioning in the membrane were performed using the PMM server [112]. Channels within *TPA* 8SβB β-barrels were analyzed using MOLEonline [29] with default settings (probe radius = 5; interior threshold = 1.1). Predicted protein-protein interfaces between *TPA* 8SβBs and host proteins were evaluated using PIZSA [34] with a Z-score cutoff of 1.8. Binding affinities (ΔG and *K*_d_) at 37°C were estimated using PRODIGY [35]. Docking analysis between TP0733 and L-ornithine (CID: 6262) was performed using CB-Dock3 [113]. Structures were visualized and analyzed using UCSF Chimera (ver. 1.16) [114]. Linear and discontinuous B-cell epitopes (BCEs) were predicted using ElliPro [47] with a threshold of 0.8. Graphical representations of BCE positions were generated using MacVector (ver. 18.8.3) (MacVector, Inc.).

### Variant calling for *TPA* 8SβB in clinical strains

Variant amino acid residues within 8SβB orthologs in *TPA* clinical strains were identified using previously published genomes [97, 115]. Sequence alignments were performed with bwa (ver. 0.7.17) [116] against the *TPA* Nichols (NC_021490.2) and SS14 (NC_021508.1) NCBI reference genomes [106]. Sequences underwent filtering post-alignment to exclude reads with excessive mismatches, excessive soft or hard clipping, chimeric reads, and/or low mapping quality. Only sequences with at least 80% coverage by at least three reads were used for subsequent analyses. Variant calling was conducted using GATK HaplotypeCaller (ver. 4.4) [117, 118]. Consensus genomes were constructed with GATK FastaAlternateReferenceMaker [117, 118]. Lineage designations for *TPA* clinical strains were assigned as previously described [94, 97]. Sequences for intact and degenerate 8SβB loci were extracted and their proteoforms analyzed using a Biopython script available at https://github.com/ebbettin/UCH_SRL. Residues under selective pressure were identified using maximum likelihood-based analysis tools available on the Datamonkey webserver [119]. Evidence of gene-wide selection was assessed using Branch-site Unrestricted Statistical Test for Episodic Diversification (BUSTED) [120]. Residues under diversifying selection were identified using Fixed Effects Likelihood (FEL) [120] with a *p*-value threshold of 0.1.

### Hybridization capture RNA-seq

Total RNA was isolated from *TPA* Nichols and SS14 reference strains harvested from rabbit testes (minimum of 3 biological replicates per strain) using ZymoBIOMICS RNA miniprep Kit (Zymo Research, Irvine, CA, USA) and treated twice with Turbo DNAse (ThermoFisher Scientific). DNA-free RNA (500 ng) was converted to cDNA using SuperScript IV reverse transcriptase (ThermoFisher Scientific) followed by second-strand synthesis with Klenow fragment (New England Biolabs, Ipswich, MA, USA). DNA concentrations were measured with the Qubit High Sensitivity double-stranded DNA kit and Qubit 2.0 Fluorometer (ThermoFisher Scientific). Libraries were prepared with the KAPA HyperCap Evolved kit (KAPA Biosystems, Inc., Wilmington, MA, USA) using custom HyperCap probes (Roche Diagnostics, Indianapolis, IN) according to the manufacturer’s instructions. Libraries were sequenced at the UConn Center for Genome Innovation (Storrs, CT, USA) on an Illumina MiSeq Micro (ver. 2) 300-cycle flow cell that generated 150-nucleotide paired-end reads. Raw reads were trimmed using Sickle (ver. 1.3.3; available from https://github.com/najoshi/sickle) and mapped using EDGE-pro (ver. 1.1.3) [121]. To enable direct comparisons between strains, genome annotations were harmonized using Liftoff to identify corresponding coding sequences and assign consistent gene identifiers across both genomes. Custom FASTA, protein translation table (PTT), and ribosomal/transfer RNA (RNT) files were generated for each strain using this unified annotation (available on request). Normalized read counts and differences in gene expression between *TPA* Nichols and SS14 reference strains were determined using DESeq2 [122]. Raw data were deposited in the NCBI Sequence Read Archive (SRA) database under Bioproject PRJNA1441567.

### Recombinant protein production

Codon-optimized synthetic genes encoding *Pyrococcus furiosus* thioredoxin (*Pf*Trx) protein scaffolds containing single extracellular loops (ECLs) and N-terminal 6xHis- and C-terminal Avi-tags [48, 123] were purchased from Integrated DNA Technologies (Coralville, IA, USA). *Pf*Trx-ECL scaffolds were biotinylated *in vivo* during expression in *E. coli* BL21 (DE3) transformed with BirA (BPS Bioscience, San Diego, CA) in Overnight Express LB autoinduction media (ThermoFisher Scientific) containing 100 μg/ml kanamycin, 50 μg/ml spectinomycin and 50 μM D-biotin as previously described [123]. Recombinant proteins were purified over Ni-NTA resin (Qiagen) followed by size-exclusion chromatography (SEC) as previously described [123].

### ELISA reactivity of *Pf*Trx-scaffolded 8SβB ECLs with immune rabbit sera

To identify ECLs recognized by antibodies in Nichols and SS14 reference strain immune rabbit sera (IRS), flat-bottom Immuno 96-well plates (ThermoFisher Scientific) were coated overnight at 4°C with streptavidin (ThermoFisher Scientific) diluted to 2 ng per ml in 0.1 M sodium bicarbonate (pH 8.5). After washing with phosphate-buffer saline containing 0.1% Tween 20 (PBST), wells were blocked for 1 h at room temperature (RT) with PBST containing 15% normal goat serum, 4% lactalbumin and 0.05% sodium azide and then coated for 1 h with biotinylated *Pf*Trx-ECL or empty scaffold (200 ng per well) diluted in the same buffer. After washing, immune rabbit sera (IRS) from animals infected with either *TPA* Nichols or SS14 [48] was added in 2-fold serial dilutions (1:20 starting dilution) in PBST containing 5% bovine serum albumin (BSA) and incubated for 1 h at RT. After washing as described above, HRP-conjugated goat anti-rabbit IgG (diluted 1:10,000 in PBST containing 15% normal goat serum and 4% lactalbumin) was added, followed by incubation for 1 h at RT. Plates were washed and developed with TMB single solution (ThermoFisher Scientific). Reactions were stopped with 0.3 M HCl after 15 min incubation and plates were read at an optical density (OD) of 450 nm. Areas under the curve (AUC) [124] for each scaffolded ECL were calculated following subtraction of values for the *Pf*Trx-empty control [48]. Graphical representations were generated using Prism (ver. 10.4.2) (GraphPad Software, San Diego, CA, USA).

## Acknowledgments

We would like to thank Arlene Sena and Chris Hennelly for their invaluable contributions to genome sequencing of *TPA* clinical strains. We also express our gratitude to study participants, as well as research, clinical, and administrative staff across clinical sites. Lastly, we thank Kristina Delgado, Carson Karanian, Mason Mansovari, Renee Pontbriant and Crystal Vicente for their assistance with assessing antibody responses to 8SβB ECLs.

The authors used an artificial intelligence language model for English language editing during manuscript preparation. However, the manuscript is original to the authors, who take responsibility for its content.

## Funding

This research project was funded by Coefficient Giving (KLH and MJC), the Gates Foundation (INV-036560 and INV-039669) and the National Institutes of Health/National Institute for Allergy and Infectious Disease (NIAID) (U19 AI144177 and U01 182179 to JDR; T32AI007151 and T32AI007001 to FA) and strategic research dollars from Connecticut Children’s (JDR, and KLH).

## Supporting information

**S1 Fig. Computational prediction of signal peptides, signal peptidase cleavage sites, and start codon sequences for *TPA* 8SβBs.** (A) Signal peptide probabilities for manually curated *TPA* 8SβB coding sequences (CDS) predicted using SignalP 6.0 (REF). Vertical dashed green lines indicate the predicted N-terminal signal peptidase cleavage sites (CS). Sec/SPI, signal peptides (SP) transported by the Sec translocon and cleaved by Signal Peptidase I (Lep). Sec/SPII, lipoprotein SP transported by the Sec translocon and cleaved by Signal Peptidase II (Lsp). n-region, N-terminal region of the SP. h-region, central hydrophobic region of SP. c-region, C-terminal region of the SP. (B) Translational start sites manually curated for six (TP0127, TP0315, TP0347, TP0479, TP0619, and TP0697) of the eleven *TPA* 8SβB proteins based on SP predictions. Start codons for coding sequences in this study differed from those annotated in the *TPA* Nichols reference strain (NCBI accession NC_021490.2) between 12 and 45 amino acid residues.

**S2 Fig. RNA-seq read mapping of the 5′ regions of *TPA* 8SβB coding sequences highlights the positions of curated start codons, putative ribosome-binding sites, and polyG tracts.** Read coverage (blue) across the 5′ coding regions of *TPA* 8SβB-encoding genes. PolyG tracts and putative ribosome-binding sites (RBS) are highlighted in brown and purple, respectively. Read mapping data support the revised translation start sites (green boxes) for six manually curated 8SβBs (A) as well as the four originally annotated start sites (B, red boxes) in the Nichols reference strain (NCBI accession NC_021490.2).

**S3 Fig. The periplasmic extension of TP0733 is predicted to be intrinsically disordered.** (A) PrDOS server predicts regions in the mature TP0733 sequence to form intrinsically disordered regions (IDR). (B) Predicted IDRs correspond to the periplasmic N-terminal region and part of ECL1 of TP0733 AlphaFold 3 model.

**S4 Fig. Hydrophobicity analysis of *TPA* 8SβBs.** (A) AlphaFold 3 structural models for *TPA* Nichols 8SβB proteins (minus signal peptides) display hydrophobic belts (yellow) characteristic of outer membrane-spanning β-barrel proteins. (B) Sectional views of models in A highlighting the hydrophilic (blue) barrel interiors of all eight full-length 8SβBs.

**S5 Fig. Structural diversity of functionally characterized 8SβBs.** Experimentally determined X-ray structures elucidate the functional versatility of prototypical 8SβBs for small-molecule transport (*E. coli* OmpW, PDB ID: 2F1V; *A. baumannii* CarO, PDB ID: 4RL9; *P. aeruginosa* OprG, PDB ID: 2X27), maintenance of outer membrane integrity (*E. coli* OmpA, PDB ID: 1BXW), host-cell adhesion (*N. gonorrhoeae* Opa60, PDB ID: 2MAF; *Y. pestis* Ail, PDB ID: 3QRA), and lipid A palmitoylation (*E. coli* PagP, PDB ID: 3GP6). Dashed lines denote the approximate position of the outer membrane.

**S6 Fig. Validation of predicted interactions between *TPA* 8SβB proteins and host ligands.** (A) AlphaFold 3 complexes between *TPA* 8SβB proteins and human CEACAM1, CEACAM5, CEACAM6, fibronectin type III repeat 9 (FNDC9), vitronectin hemopexin domain (VTN), and complement factor H sushi domains 6-7 (FH) were submitted to PIZSA to assess the biological likelihood and stability of the resulting protein-protein interfaces. (B) Complexes with *Z*-scores >1.8 were considered high-confidence and subsequently analyzed. DNB, does not bind.

**S7 Fig. PolyG tracts are widely distributed across 8SβB loci.** Alignment of 8SβB coding regions from 1706 clinical strains with Nichols and SS14 reference sequences revealed a total of 10 upstream and 21 intragenic PolyG tracts within 8SβB loci. Tracts upstream of DUF2715 subgroup I orthologs (*tp0127*, tp0315*, tp0618*,* and *tp0619*) were invariant (yellow dashes), while PolyGs upstream *tp0126, tp0347, tp0479,* and *tp0697* exhibited length polymorphisms (red dashes) among clinical strains. In contrast, variable intragenic polyG tracts were observed exclusively within truncated DUF2715 subgroup I orthologs *(tp0127*, tp0315*, and TP0618*)*.

**S8 Fig. Variable PolyG tracts in *TPA* reference and clinical strains.** Variable PolyG tracts were identified upstream four (A), and within two (B) 8SβB-encoding genes. Alleles found in Nichols and SS14 reference strains are boxed. Asterisks in B denote the alleles corresponding to full-length proteoforms.

**S9 Fig. *TPA* 8SβB proteins regain their predicted β-barrel topologies following *in silico* correction of frameshift mutations.** (A) The restored proteins exhibited high sequence and structural similarity to TP0619, a member of the DUF2715 subgroup I family. (B) Restoration of the open reading frames for TP0127*, TP0315* (identical to TP0619), and TP0618* enabled the prediction of high-confidence β-barrel structures by AlphaFold 3. (C) Sequence alignment of DUF2715 subgroup I orthologs shows that differences are concentrated within ECLs. (D) Alphafold 3 complexes for the restored DUF2715 subgroup I proteins TP0127* and TP0618* did not displayed high-confidence (*Z*-score <1.8) or exhibited substantially weaker predicted binding free energies (ΔG, kcal/mol) and dissociation constants (Kd) for interactions with VTN and VE-cad when compared to TP0619/TP0315*.

**S1 Table. Description of 8SβB gene neighborhoods in the *TPA* Nichols reference genome (NC_021490.2)**

**S2 Table. Linear and discontinuous B-cell epitopes predicted using ElliPro.**

**S3 Table. Metadata associated with the 1706 clinical *TPA* strains used in this study.**

**S4 Table. *TPA* 8SβB are highly conserved across clinical strains isolated from patients with primary or secondary syphilis.**

**S5 Table. Description of codon variants within 1706 analyzed *TPA* clinical strains.**

**S6 Table. Analysis of 8SβB repertoires within the 1706 analyzed *TPA* clinical strains.**

## Notes

### Competing Interest Statement

The authors have declared no competing interest.

