## Supplementary Figures for "*Treponema pallidum* eight-stranded beta-barrels: an evolving, versatile family of outer membrane proteins in the service of stealth pathogenicity"

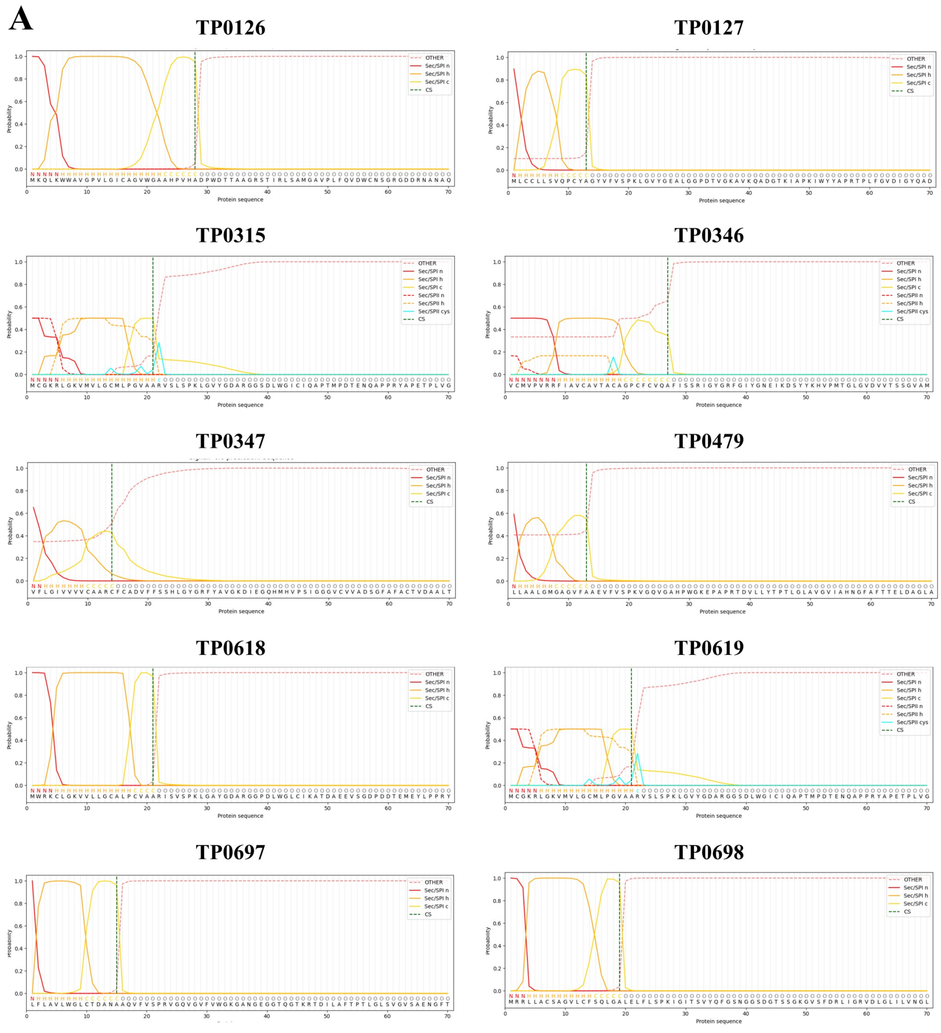


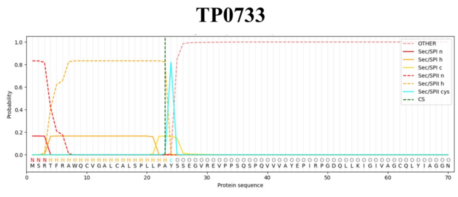


**B**

**Start codon adjustments and Signal peptide predictions**

| Protein | Size difference (aa) | Start position in NC_021490.2 (bp) | |
|---|---|---|---|
| | | Annotated | This study |
| TP0127 | -12 | 149809 | 149845 |
| TP0315 | -35 | 332305 | 332200 |
| TP0347 | -45 | 373110 | 373245 |
| TP0479 | -45 | 511161 | 511026 |
| TP0619 | -35 | 672662 | 672557 |
| TP0697 | -22 | 767373 | 767307 |

**S1 Fig. Computational prediction of signal peptides, signal peptidase cleavage sites, and start codon sequences for *TPA* 8SβBs.** (A) Signal peptide probabilities for manually curated *TPA* 8SβB coding sequences (CDS) predicted using SignalP 6.0 (REF). Vertical dashed green lines indicate the predicted N-terminal signal peptidase cleavage sites (CS). Sec/SPI, signal peptides (SP) transported by the Sec translocon and cleaved by Signal Peptidase I (Lep). Sec/SPII, lipoprotein SP transported by the Sec translocon and cleaved by Signal Peptidase II (Lsp). n-region, N-terminal region of the SP. h-region, central hydrophobic region of SP. c-region, C-terminal region of the SP. (B) Translational start sites manually curated for six (TP0127, TP0315, TP0347, TP0479, TP0619, and TP0697) of the eleven *TPA* 8SβB proteins based on SP predictions. Start codons for coding sequences in this study differed from those annotated in the *TPA* Nichols reference strain (GenBank accession NC_021490.2) between 12 and 45 amino acid residues.

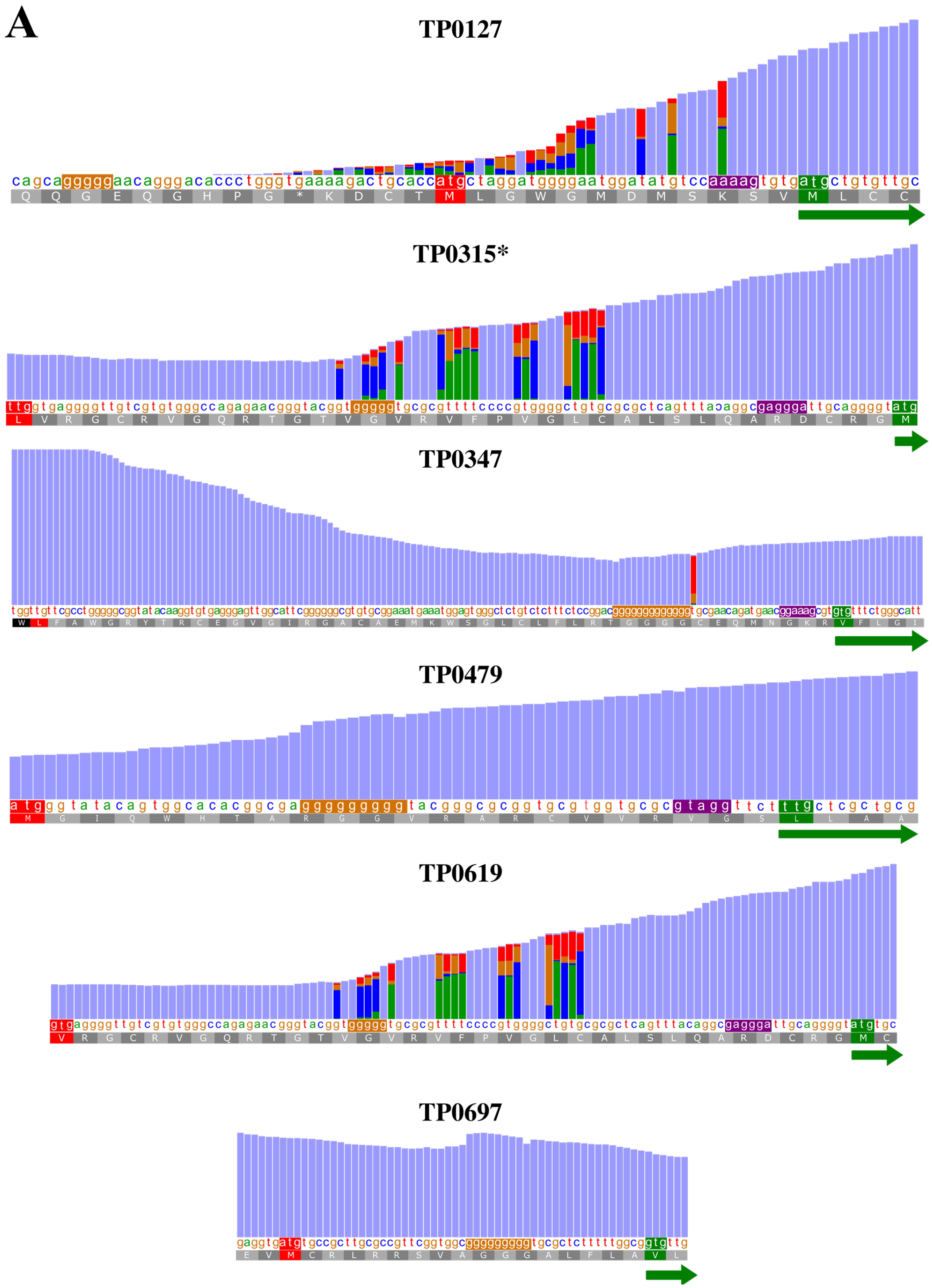
A
TP0127
TP0315*
TP0347
TP0479
TP0619
TP0697

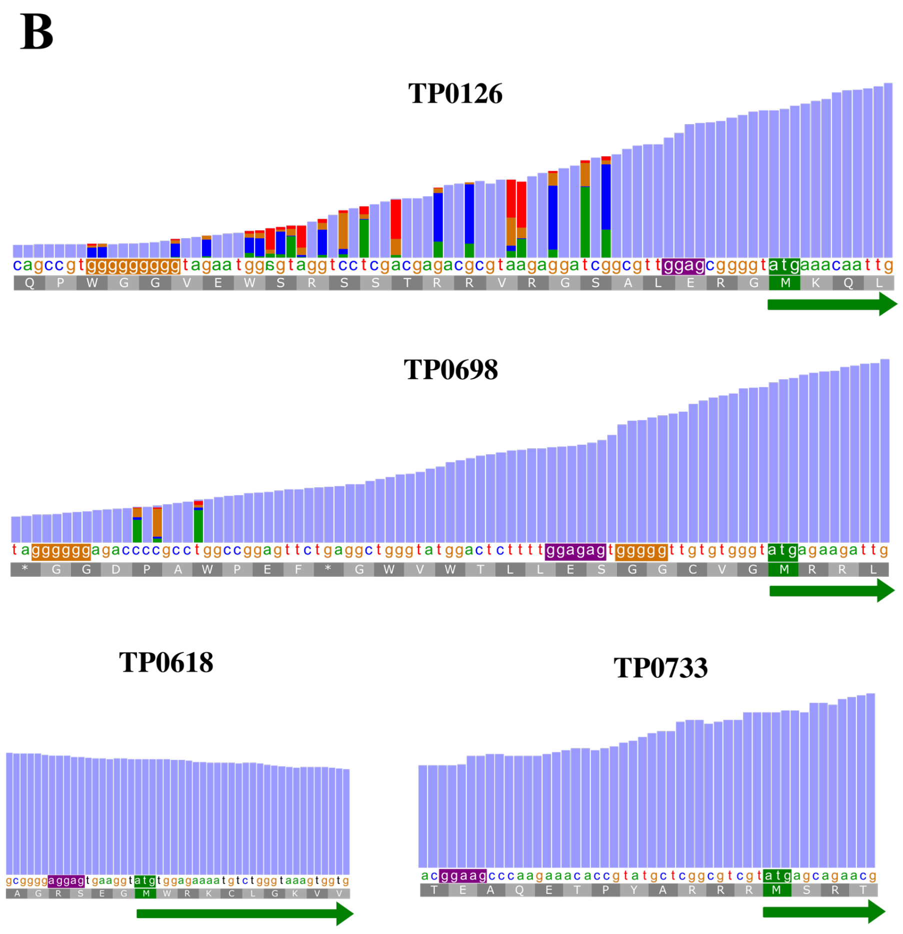


**S2 Fig. RNA-seq read mapping of the 5′ regions of *TPA* 8SβB coding sequences highlights the positions of curated start codons, putative ribosome-binding sites, and polyG tracts.** Read coverage (blue) across the 5′ coding regions of *TPA* 8SβB-encoding genes. PolyG tracts and putative ribosome-binding sites (RBS) are highlighted in brown and purple, respectively. Read mapping data support the revised translation start sites (green boxes) for six manually curated 8SβBs (A) as well as the four originally annotated start sites (B, red boxes) in the Nichols reference strain (GenBank accession NC_021490.2).

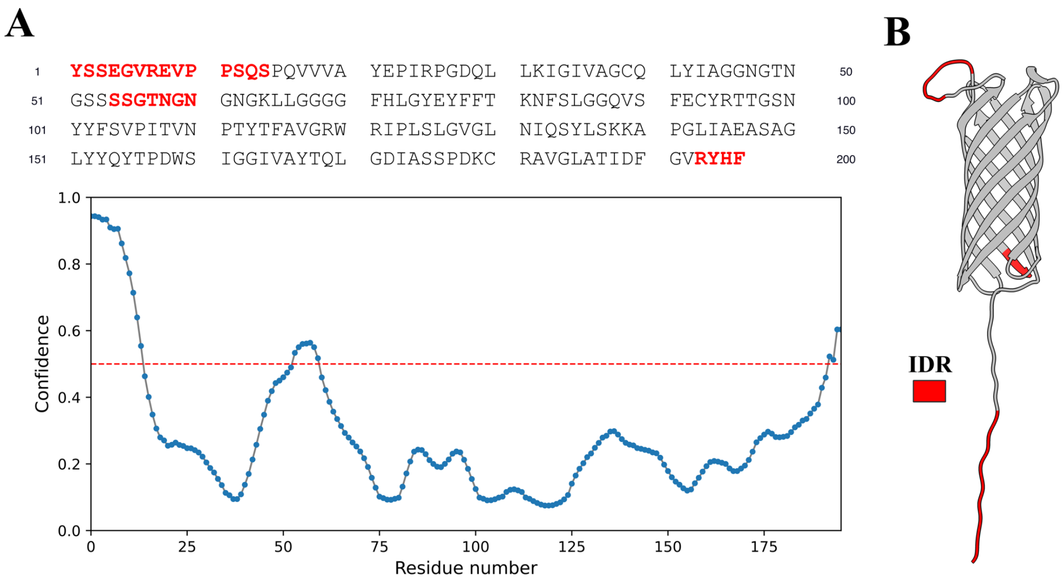


**S3 Fig. The periplasmic extension of TP0733 is predicted to be intrinsically disordered.** (A) PrDOS server predicts regions in the mature TP0733 sequence to form intrinsically disordered regions (IDR). (B) Predicted IDRs correspond to the periplasmic N-terminal region and part of ECL1 of TP0733 AF3 model.

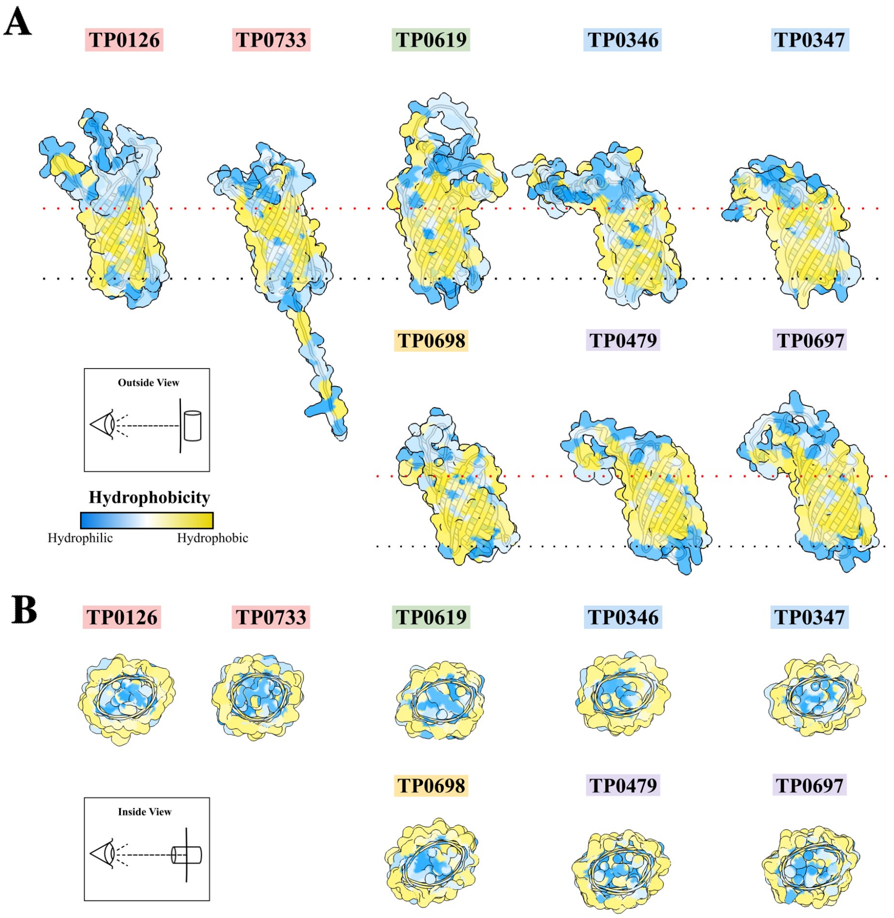


**S4 Fig. Hydrophobicity analysis of TPA 8SβBs.** (A) AlphaFold 3 structural models for TPA Nichols 8SβB proteins (minus signal peptides) display hydrophobic belts (yellow) characteristic of outer membrane-spanning β-barrel proteins. (B) Sectional views of models in A highlighting the hydrophilic (blue) barrel interiors of all eight full-length 8SβBs.

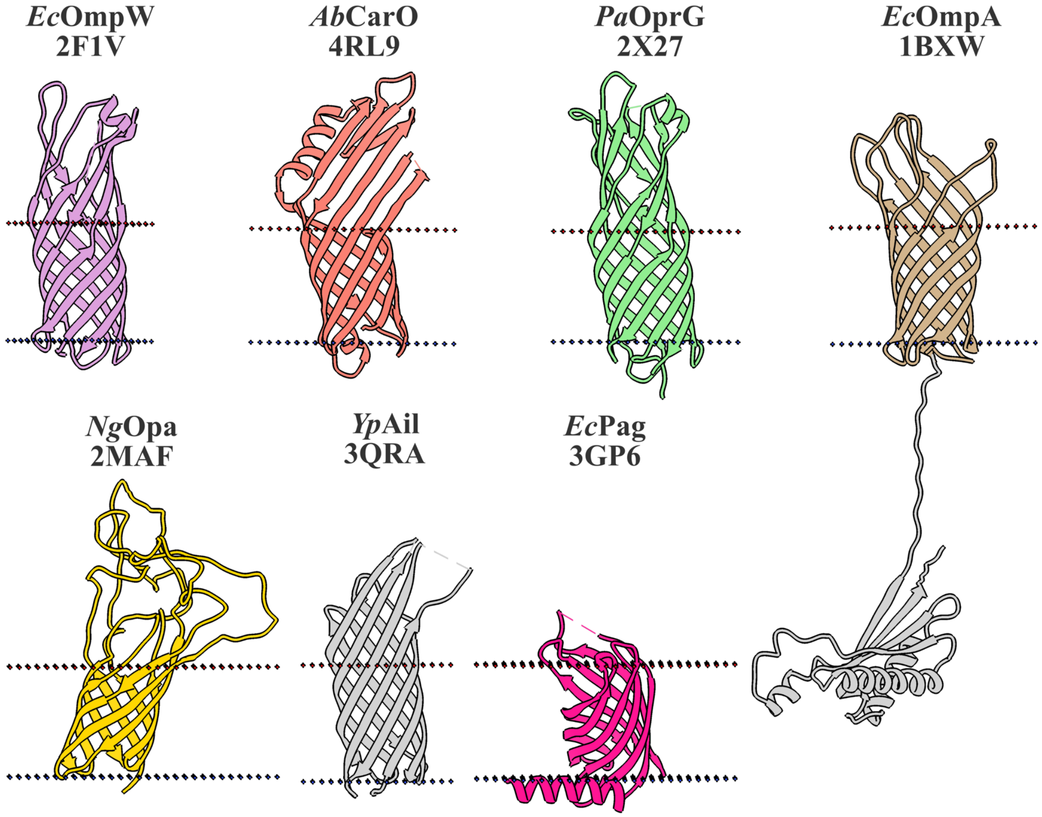


**S5 Fig. Structural diversity of functionally characterized 8SβBs.** Experimentally determined X-ray structures elucidate the functional versatility of prototypical 8SβBs for small-molecule transport (E. coli OmpW, PDB ID: 2F1V; A. baumannii CarO, PDB ID: 4RL9; P. aeruginosa OprG, PDB ID: 2X27), maintenance of outer membrane integrity (E. coli OmpA, PDB ID: 1BXW), host-cell adhesion (N. gonorrhoeae Opa, PDB ID: 2MAF; Y. pestis Ail, PDB ID: 3QRA), and lipid A palmitoylation (E. coli PagP, PDB ID: 3GP6). Dashed lines denote the approximate position of the outer membrane.

**A**

| | Z-Score | | | | | | |
|---|---|---|---|---|---|---|---|
| | **CEACAM1** | **CEACAM5** | **CEACAM6** | **VTN** | **FH** | **E-Cad** | **VE-Cad** |
| ***Ng* OPA** | 1.861 | 2.333 | 2.59 | 1.328 | DNB | 1.38 | 1.39 |
| ***Yp* Ail** | 1.469 | 1.252 | 1.238 | 1.889 | 1.927 | 0.598 | 1.46 |
| ***Ec* OmpA** | 1.673 | 0.695 | 1.378 | 0.427 | DNB | 0.53 | -1.17 |
| **TP0126** | 1.25 | 1.42 | 1.52 | 1.84 | 2.605 | 2.005 | 1.548 |
| **TP0733** | 1.06 | DNB | 0.99 | 1.32 | DNB | 1.61 | DNB |
| **TP0619** | 1.48 | 1.222 | 0.46 | 2.15 | 1.612 | -1.314 | 1.934 |
| **TP0698** | 1.46 | 1.25 | 1.25 | 1.68 | 1.831 | 0.459 | 1.565 |
| **TP0346** | 2.01 | 2.02 | 1.82 | 1.99 | 1.549 | 1.135 | 2.038 |
| **TP0347** | 2.59 | 1.84 | 2.28 | 1.61 | 1.796 | 1.158 | 0.776 |
| **TP0479** | 1.95 | 1.85 | 2.39 | 1.75 | 1.289 | 0.453 | 1.348 |
| **TP0697** | 1.93 | 1.90 | 2.43 | 2.21 | 0.879 | 1.168 | 1.988 |

**B**

**CEACAMs**

| Complex | ΔG | Kd | charged-charged | charged-polar | charged-apolar | polar-polar | polar-apolar | apolar-apolar | Total |
|---|---|---|---|---|---|---|---|---|---|
| TP0346_CEACAM1 | -9 | 4.30E-07 | 4 | 22 | 15 | 6 | 17 | 17 | 81 |
| TP0347_CEACAM1 | -7.3 | 6.60E-06 | 1 | 7 | 10 | 10 | 17 | 22 | 67 |
| TP0479_CEACAM1 | -9.6 | 1.70E-07 | 7 | 15 | 22 | 5 | 17 | 27 | 93 |
| TP0697_CEACAM1 | -6.2 | 4.10E-05 | 2 | 6 | 8 | 7 | 13 | 10 | 46 |
| TP0346_CEACAM5 | -7.7 | 3.60E-06 | 10 | 15 | 14 | 4 | 10 | 20 | 73 |
| TP0347_CEACAM5 | -7.3 | 7.10E-06 | 2 | 11 | 9 | 5 | 15 | 20 | 62 |
| TP0479_CEACAM5 | -10.4 | 4.60E-08 | 11 | 14 | 24 | 6 | 21 | 27 | 103 |
| TP0697_CEACAM5 | -9.7 | 1.50E-07 | 6 | 16 | 20 | 7 | 24 | 28 | 101 |
| TP0346_CEACAM6 | -4.8 | 3.80E-04 | 0 | 3 | 7 | 11 | 9 | 19 | 49 |
| TP0347_CEACAM6 | -8.4 | 1.20E-06 | 0 | 6 | 8 | 4 | 19 | 19 | 56 |
| TP0479_CEACAM6 | -10.4 | 4.90E-08 | 4 | 12 | 20 | 3 | 21 | 35 | 95 |
| TP0697_CEACAM6 | -7.5 | 5.20E-06 | 1 | 2 | 11 | 5 | 16 | 18 | 53 |

**Vitronectin**

| Complex | ΔG | Kd | charged-charged | charged-polar | charged-apolar | polar-polar | polar-apolar | apolar-apolar | Total |
|---|---|---|---|---|---|---|---|---|---|
| TP0126_VTN | -9.3 | 2.90E-07 | 12 | 16 | 21 | 12 | 16 | 6 | 83 |
| TP0346_VTN | -10.8 | 2.20E-08 | 12 | 7 | 23 | 0 | 13 | 27 | 82 |
| TP0619_VTN | -20.6 | 3.00E-15 | 17 | 21 | 68 | 4 | 38 | 81 | 229 |
| TP0697_VTN | -11.2 | 1.4E-08 | 10 | 17 | 26 | 2 | 21 | 13 | 89 |

**Factor-H**

| Complex | ΔG | Kd | charged-charged | charged-polar | charged-apolar | polar-polar | polar-apolar | apolar-apolar | Total |
|---|---|---|---|---|---|---|---|---|---|
| TP0126_FH | -7.4 | 6.40E-06 | 5 | 1 | 10 | 0 | 2 | 6 | 18 |
| TP0698_FH | -11.9 | 3.90E-09 | 7 | 15 | 24 | 3 | 24 | 23 | 73 |

**E-Cadherin**

| Complex | ΔG | Kd | charged-charged | charged-polar | charged-apolar | polar-polar | polar-apolar | apolar-apolar | Total |
|---|---|---|---|---|---|---|---|---|---|
| TP0126_E-Cad | -11.9 | 4.10E-09 | 7 | 12 | 18 | 6 | 27 | 15 | 70 |

**VE-Cadherin**

| Complex | ΔG | Kd | charged-charged | charged-polar | charged-apolar | polar-polar | polar-apolar | apolar-apolar | Total |
|---|---|---|---|---|---|---|---|---|---|
| TP0619_VE-Cad | -8.7 | 7.90E-07 | 9 | 14 | 6 | 6 | 17 | 7 | 52 |
| TP0697_VE-Cad | -6.9 | 1.30E-05 | 3 | 6 | 10 | 11 | 16 | 10 | 46 |

**S6 Fig. Validation of predicted interactions between *TPA* 8SβB proteins and host ligands.** AlphaFold 3 complexes between *TPA* 8SβB proteins and human CEACAM1, CEACAM5, CEACAM6, fibronectin type III repeat 9 (FNDC9), vitronectin hemopexin domain (VTN), and complement factor H sushi domains 6-7 (FH) were submitted to PIZSA to assess the biological likelihood and stability of the resulting protein-protein interfaces. (B) Complexes with *Z*-scores >1.8 were considered high-confidence and subsequently analyzed. DNB. Does not bind.

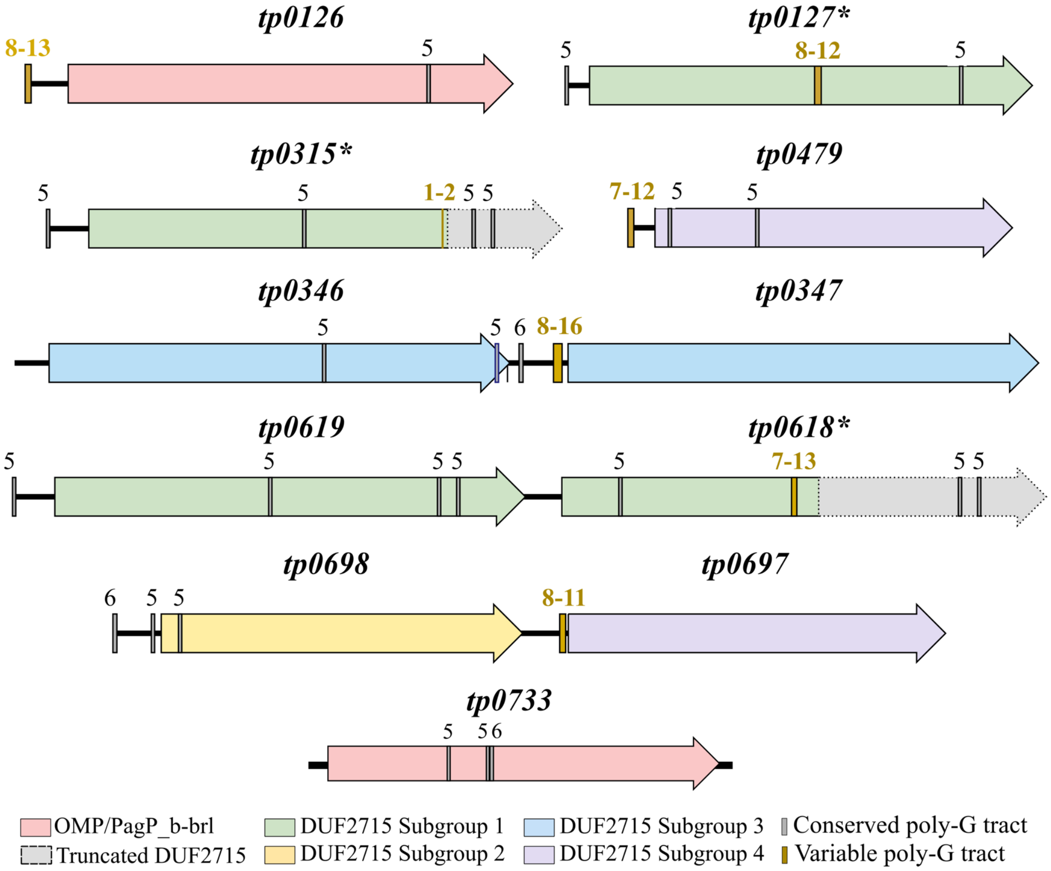


**S7 Fig. PolyG tracts are widely distributed across 8SβB loci.** Alignment of 8SβB coding regions from 1706 clinical strains with Nichols and SS14 reference sequences revealed a total of 10 upstream and 21 intragenic PolyG tracts within 8SβB loci. Tracts upstream from DUF2715 subgroup I orthologs (tp0127*, tp0315*, tp0618*, and tp0619) were invariant (yellow dashes), while PolyGs upstream tp0126, tp0347, tp0479, and tp0697 exhibited length polymorphisms (red dashes) among clinical strains. In contrast, variable intragenic polyG tracts were observed exclusively within truncated DUF2715 subgroup I orthologs (tp0127*, tp0315*, and TP0618*).

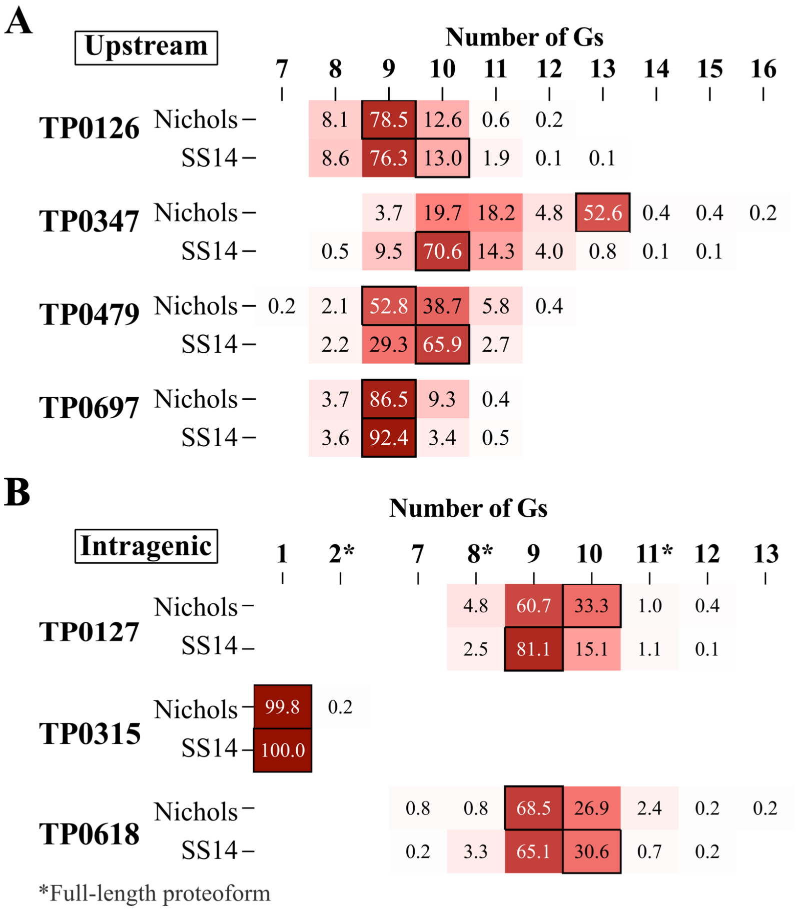


**S8 Fig. Variable PolyG tracts in TPA reference and clinical strains.** Variable PolyG tracts were identified upstream four (A), and within two (B) 8SβB-encoding genes. Alleles found in Nichols and SS14 reference strains are boxed. Asterisks in B denote the alleles corresponding to full-length proteoforms.

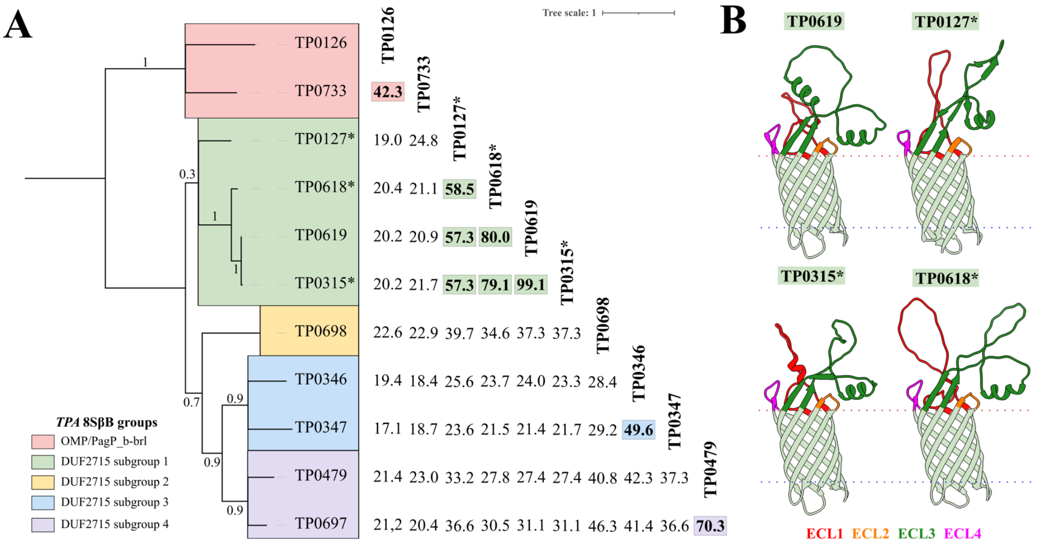


C

**Similarity Matrix (%)**

| | TP0127 | TP0618 (Barrel) | TP0315 (Barrel) | TP0619 (Barrel) |
|---|---|---|---|---|
| **TP0127** | | 76.1 | 75 | 75 |
| **TP0618** | 38.2 | | 92.5 | 94.4 |
| **TP0315** | 36.1 | 67.2 | | 98.1 |
| **TP0619** | 36.1 | 67.2 | 100 | |
| | ECLs | ECLs | ECLs | |

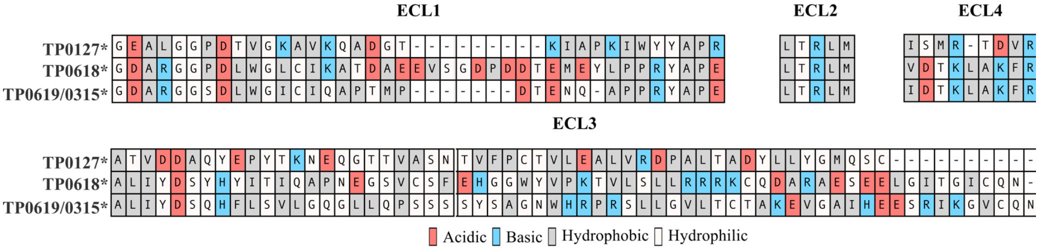


D

| | Vitronectin Z-Score | Vitronectin ΔG | Vitronectin Kd |
|---|---|---|---|
| **TP0619** | 2.15 | -20.6 | 3.00E-15 |
| **TP0127** | 1.493 | | |
| **TP0618** | 1.602 | | |

| | VE-cad Z-Score | VE-cad ΔG | VE-cad Kd |
|---|---|---|---|
| **TP0619** | 1.934 | -8.7 | 7.90E-07 |
| **TP0127** | 1.428 | | |
| **TP0618** | 2.474 | -6.3 | 3.40E-05 |

**S9 Fig. TPA 8SβB proteins regain their predicted β-barrel topologies following in silico correction of frameshift mutations.** (A) The restored proteins exhibited high sequence and structural similarity to TP0619, a member of the DUF2715 subgroup I family. (B) Restoration of the open reading frames for TP0127*, TP0315* (identical to TP0619), and TP0618* enabled the prediction of high-confidence β-barrel structures by AlphaFold 3. (C) Sequence alignment of DUF2715 subgroup I orthologs shows that differences are concentrated within ECLs. (D) Alphafold 3 complexes for the restored DUF2715 subgroup I proteins TP0127* and TP0618* did not displayed high-confidence (*Z*-score <1.8) or exhibited substantially weaker predicted binding free energies (ΔG, kcal/mol) and dissociation constants (Kd) for interactions with VTN and VE-cad when compared to TP0619/TP0315*.
