## Supplementary Tables for "*Treponema pallidum* eight-stranded beta-barrels: an evolving, versatile family of outer membrane proteins in the service of stealth pathogenicity": S4 Table.docx

| S4 Table. *TPA* 8SβB are highly conserved across clinical strains isolated from patients with primary or secondary syphilis. | | | | | |
| --- | --- | --- | --- | --- | --- |
|  | **Mutation** | **Mapping** | **Count** | | **%** |
|  |  |  | **Nichols** | **SS14** |  |
| TP0126 | G62E | ECL1 | 2 |  | 0.11 |
| TP0346 | G36R | ECL1 |  | 1 | 0.06 |
|  | R37H | ECL1 | 2 | 1222 | 71.74 |
|  | R37Y | ECL1 | 3 |  | 0.18 |
|  | Y50C | ECL1 |  | 1 | 0.06 |
|  | A103V | β-strand 5 | 2 |  | 0.12 |
|  | N149S | ECL4 | 1 |  | 0.06 |
|  | M154V | ECL4 | 1 | 1223 | 71.74 |
|  | K163E | ECL4 | 100 |  | 5.80 |
|  | T167M | ECL4 | 1 |  | 0.06 |
|  | S187P | ECL4 |  | 1 | 0.06 |
|  | G193R | ECL4 |  | 1 | 0.06 |
|  | G193E | ECL4 |  | 1 | 0.06 |
| TP0347 | G86D | β-strand 4 | 1 |  | 0.08 |
|  | A93E | Turn 2 | 1 |  | 0.08 |
| TP0479 | D128N | β-strand 7 | 7 |  | 0.41 |
| TP0619 | C131R | β-strand 4 |  | 1223 | 71.68 |
|  | R133Q | β-strand 4 |  | 1223 | 71.68 |
| TP0697 | I50V | ECL1 |  | 1 | 0.06 |
|  | V61M | β-strand 2 | 7 |  | 0.41 |
|  | A150E | ECL4 |  | 1 | 0.06 |
| TP0733 | A43V | Periplasmic |  | 4 | 0.23 |
|  | G69E | ECL1 | 5 |  | 0.29 |
|  | N73D | ECL1 | 61 |  | 3.53 |
|  | G74D | ECL1 |  | 1 | 0.06 |
|  | G74S | ECL1 |  | 2 | 0.12 |
|  | S75P | ECL1 |  | 1 | 0.06 |
|  | S76P | ECL1 | 7 |  | 0.41 |
|  | S78F | ECL1 | 4 |  | 0.23 |
|  | G79S | ECL1 | 26 |  | 1.53 |
|  | G74_T80del | ECL1 |  | 1 | 0.06 |
|  | T179N | Turn 3 |  | 12 | 0.70 |
|  | D195E | ECL4 |  | 1 | 0.06 |
|  | S198N | ECL4 | 1 |  | 0.06 |
|  | S199I | ECL4 |  | 1 | 0.06 |
|  | S199P | ECL4 | 37 | 5 | 2.43 |
|  | P200T | ECL4 |  | 2 | 0.12 |
|  | P200L | ECL4 |  | 1 | 0.06 |
|  | C203R | ECL4 |  | 8 | 0.46 |
|  | R204G | ECL4 |  | 2 | 0.12 |
